# Active pursuit gates egocentric coding in the retrosplenial cortex

**DOI:** 10.64898/2026.02.14.705560

**Authors:** Pearl Saldanha, Martin Bjerke, Benjamin A. Dunn, Jonathan R. Whitlock

## Abstract

Spatial navigation is commonly studied in static environments, but adaptive behavior frequently hinges on tracking moving goals in real time. Active pursuit exemplifies this challenge: it is an inherently egocentric spatial behavior requiring continuous localization of a moving target, yet the neural coding schemes supporting it remain poorly understood. We therefore performed Neuropixels recordings in the retrosplenial cortex (RSC), a hub for egocentric-to-allocentric reference frame transformations, during naturalistic bait-chasing. We identified head-centered target-encoding neurons that were functionally distinct from cells encoding environmental boundaries or static objects. Whereas egocentric boundary encoding was task-invariant, target-coding cells were specific to chasing and exhibited task-dependent retuning, characterized by enhanced ego-centric representation and reduced allocentric head-direction signaling during pursuit. Together, these findings demonstrate that classical egocentric and allocentric codes coexist with novel target tuning in the RSC, where target-coding neurons dynamically reweight their reference frames according to behavioral demands during pursuit.

## INTRODUCTION

The retrosplenial cortex (RSC) sits at a crossroads of spatial navigation systems in the brain, serving to translate between allocentric and egocentric reference frames [1, 2, 3, 4, 5, 6, 7, 8, 9, 10, 11, 12, 13, 14]. This view has emerged from decades of work demonstrating that RSC neurons encode a rich repertoire of spatial and self-motion variables, including allocentric head direction [12, 15], spatial position [16, 17], rotational head velocity [18], visual motion and landmark information [19, 20, 21], as well as the position of environmental boundaries relative to the individual [7, 8, 22]. However, these insights derive largely from studies in static or minimally structured environments, typically during open field exploration or simple navigation tasks [23, 24, 7, 8, 25]. This approach assumes that spatial mapping is optimized for representing stable layouts and does not account for more dynamic ethological behaviors. Consequently, little is known of how the RSC supports more rapid forms of navigation that require online updating of spatial information [26, 27], such as during pursuit and interception of a moving target.

This gap is significant because real-world behavior extends far beyond exploration of open spaces. Animals must not only be able to self-localize in an environment, but also track moving goals around them, adjust their actions, and flexibly switch behavioral strategies in real time.

The RSC is well positioned to manage these demands, given its prominent coding of ego-centric movement and boundaries [12, 7, 8, 22], as well as visual motion, allocentric space, landmarks and head direction [18, 28, 6]. Anatomically, the RSC receives convergent signals from visual, parietal and hippocampal areas while projecting to collicular, motor and premotor cortices [29, 30, 31], placing it to efficiently transform extrinsic signals from the world into body-centered action commands[32, 33, 10, 9, 2, 34]. Yet how this polyphony of ego- and allocentric signals is coordinated while an animal pursues a moving target is unknown, and it remains unclear whether moving targets and the environment are encoded in a common framework or by functionally specialized modules.

We addressed these questions using a bait-chasing task in which rats pursue a dynamically moving food reward. In contrast with foraging in a static arena, the pursuit task requires continual updating of the target location while both it and the animal move through space. We performed Neuropixels recordings in the RSC during this naturalistic behavior, which revealed a previously undescribed sub-class of neurons encoding the moving target relative to the head: egocentric target cells (ETCs). These neurons were distinct from egocentric boundary cells (EBCs)– they did not emerge from the EBC population, nor did they encode static objects in the arena. At the population level, ETC activity occupied a task-specific state space during pursuit, while individual ETCs exhibited task-dependent ‘remapping’, shifting from prevalent coding of allocentric head direction during foraging to encoding target location and running speed during chasing. These findings demonstrate that RSC contains modular egocentric representations for stable environmental structure and moving goals and, beyond transforming reference frames, it prioritizes them dynamically depending on the demands of behavior.

## RESULTS

### Egocentric target coding emerges in retrosplenial cortex during pursuit

To characterize spatial and egocentric tuning in the retrosplenial cortex, we recorded neural activity from four adult female Long Evans rats (3–4 months of age). Neuropixels 1.0 probes were chronically implanted in the dysgranular and granular RSC, yielding a total of *N* = 459 well-isolated single units across all animals and sessions (Figure S1). Animals were tracked with a marker-based 3D motion capture system (STAR Methods) in an open arena (140 *×* 140 *×* 50 cm) with vinyl flooring and a cue card on the north wall (Figure 1A). The recording paradigm centered on a dynamic bait-chasing task in which animals tracked a dynamically moving food reward over multiple trials that were interleaved by pauses for food consumption (Figure 1A; Video S1). The other tasks consisted of spontaneous foraging in the same arena either while it was either empty or contained a static object (10 *×* 50 cm striped cylinder).

**Figure 1:**
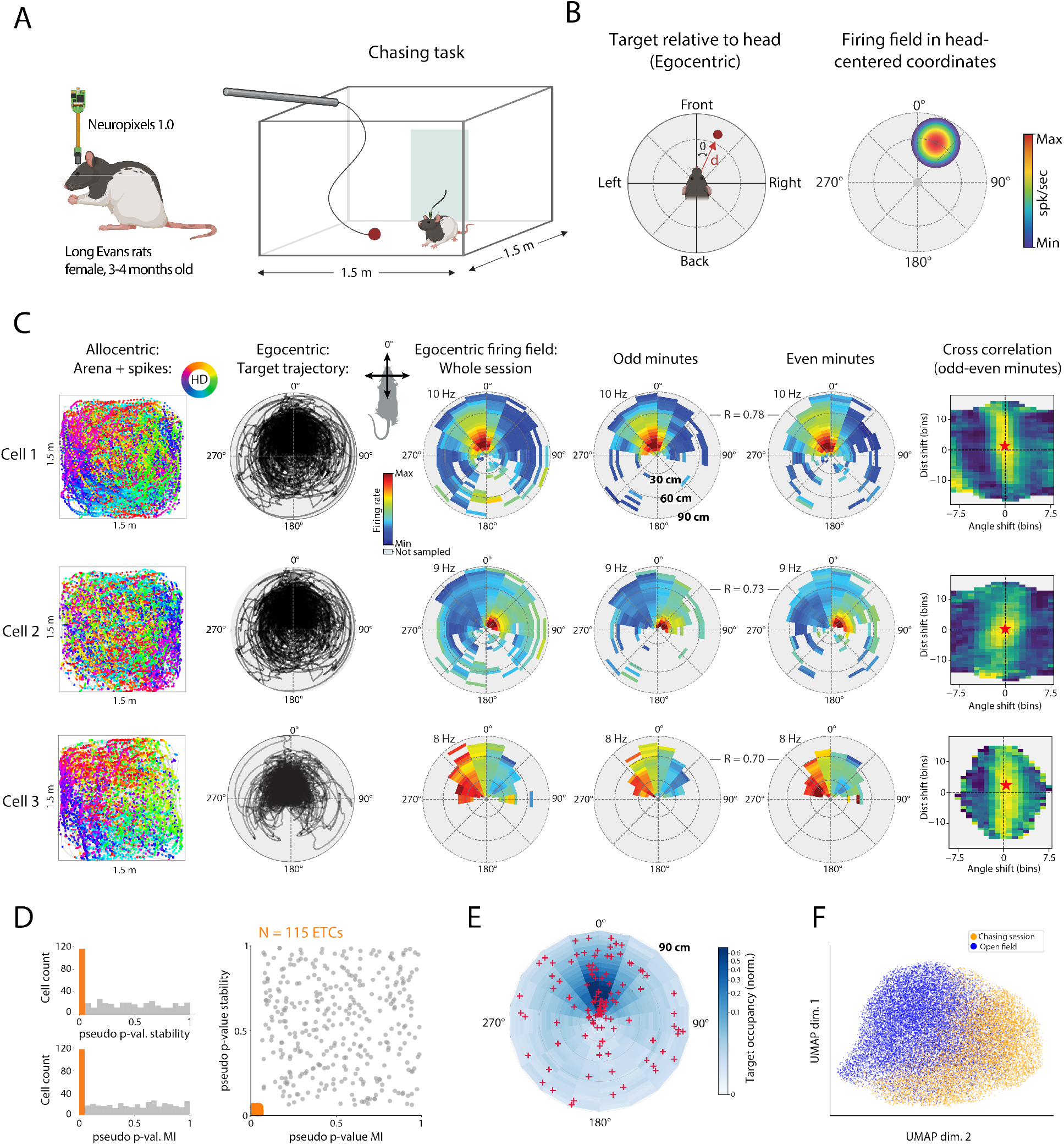
Egocentric target coding emerges in retrosplenial cortex during pursuit. (A) Schematic of the electrophysiological approach and chasing task. Adult female rats (Long-Evans, 3–4 months old) were implanted chronically with Neuropixels 1.0 probes in the RSC and. The animals moved freely in a 1.5 m *×* 1.5 m arena and chased a food bait (denoted by red dot) attached to a rod the experimenter held and moved. (B) Schematic for generating egocentric target rate maps. Left: the target position is expressed as the angle *θ* and distance *d* from the head of the animal. Right, schematic egocentric target rate map showing the color-coded firing rate of a cell coding for the target at that specific angle and distance (up to 90 cm, indicated by rings). (C) Examples of three egocentric target cells (ETCs). Each row depicts (left to right): the path of the animal in the area with spike positions (colored by head direction) overlaid; egocentric target occupancy; rate map showing the preferred target location of the cell relative to the head (whole session average); rate maps with data from odd or even minutes only; cross-correlogram between odd and even minute maps showing the angle (x-axis) and distance (y-axis) of the shift of the correlation peak (red star). (D) ETC classification summaries. Left, histogram of pseudo-*p* values for within-session (even-odd minute) stability of the egocentric rate maps; orange bars indicate neurons exceeding the 99th percentile of the shuffled distribution. Middle, histogram of pseudo-*p* values for mutual information (MI); neurons above 99th percentile of the shuffled data shown in orange. Right, joint distribution of stability and MI pseudo-*p* values; orange points indicate neurons classified as ETCs (pseudo-*p <* 0.01 for both criteria). (E) Polar chart showing the distribution of tuning peaks around the animals’ head (red crosses) and normalized cumulative occupancy of the pursuit target relative to the head (blue shading); both summed across all four animals. (F) UMAP embedding of ETC population activity. Each point represents a single time bin, colored by behavioral context (blue: open field; orange: chasing). The population activity occupied distinct regions of state space across contexts (STAR Methods), indicating robust differences in population dynamics between the two session types.

We first examined whether individual RSC neurons encoded the egocentric position of the bait in the chasing task, independent of its allocentric location. To do so, we constructed egocentric target rate maps (ETRs) by expressing the bait’s position in a head-centered polar coordinate frame directionally aligned to the animal’s nose (Figure 1B). Egocentric angle (*θ*) spanned 0^◦^–360^◦^ in 20^◦^ bins, and distance (*d*) was binned in 4.5 cm increments up to 90 cm. For each neuron, the firing rate per bin was calculated as the total spike count divided by occupancy, yielding a head-centered representation of target-related spiking activity (Figure 1B, right). This revealed firing fields that stably encoded the distance and direction of the bait relative to the head (Figure 1C; Video S2).

Neurons were classified as egocentric target cells (ETCs) based on two metrics: the Skaggs mutual information (MI) index, which measures how much information a neuron’s firing conveys about the target’s egocentric position [35], and tuning stability, measured by the Pearson correlation between alternating 30 s (even-odd) blocks. Statistical significance for both metrics was assessed using 1000 circular spike-time shuffles, with observed scores converted to pseudo-*p* values to provide a standardized (0 to 1 scale) comparison across neurons with differing null-distribution scales (Figures 1D and S2; STAR Methods). Only neurons exceeding the 99th percentile (pseudo-*p <* 0.01) of both null distributions were labeled as ETCs (Figure 1D). This conservative threshold identified 115 of 459 neurons (25.1%) as ETCs, confirming the expression of head-centered representations of the moving target comparable to those described in the posterior parietal cortex (PPC) [36]. There was no evident spatial organization or contralateral bias for the firing fields around the head (Figure 1E), and ETC firing fields were absent in open field sessions when there was no bait (Figure S3).

Having established ETCs at the single-cell level, we investigated whether active pursuit drove task-specific reorganization of their population activity. We visualized population activity using Uniform Manifold Approximation and Projection (UMAP)[37] to compare the state spaces occupied by ETCs during the chasing task versus open field foraging (Figure 1F). In the embedding, each point represents a single time bin colored by behavioral context, revealing a clear separation of population dynamics across the two tasks. We quantified the organization of the population states using complementary measures of clustering and stability. Geometric metrics confirmed the separation across contexts (Silhouette score = 0.26; Dunn index = 0.15; STAR Methods), and low (12.1%) overlap between tasks demonstrated strong state-specificity in the neural activity. Moreover, a linear discriminant analysis (LDA) classified the correct behavioral context with 91.3% accuracy (STAR Methods). Together, these results demonstrate that egocentric target coding is robust in a subset of RSC neurons and that active pursuit reconfigures their population dynamics into different regimes.

### Egocentric boundary coding is unaffected by pursuit

We next investigated whether pursuit-related activity emerged from a reassignment of the RSC’s well-characterized egocentric boundary cells (EBCs) [7, 8]. If pre-existing boundary representations were re-tuned to track salient moving targets, ETCs should appear from from the existing EBC population rather than *de novo*. We therefore tested whether egocentric boundary tuning was preserved or reconfigured when animals transitioned from open field exploration to the pursuit task.

First, we constructed egocentric boundary rate maps (EBRs) by expressing the angle and distance to the nearest arena wall in head-centered coordinates (Figure 2A). Boundary angles binned in 10^◦^ increments and distances up to 60 cm in 5 cm steps. Tuning strength was quantified using the mean resultant length (MRL; STAR Methods), following established EBC classification protocols [7]. As with ETCs, neurons were classified as EBCs only if their MRL and stability measures both exceeded the 99th percentile of the null distribution generated by 1,000 circular spike-time shuffles (STAR Methods) (Figure 2B).

**Figure 2:**
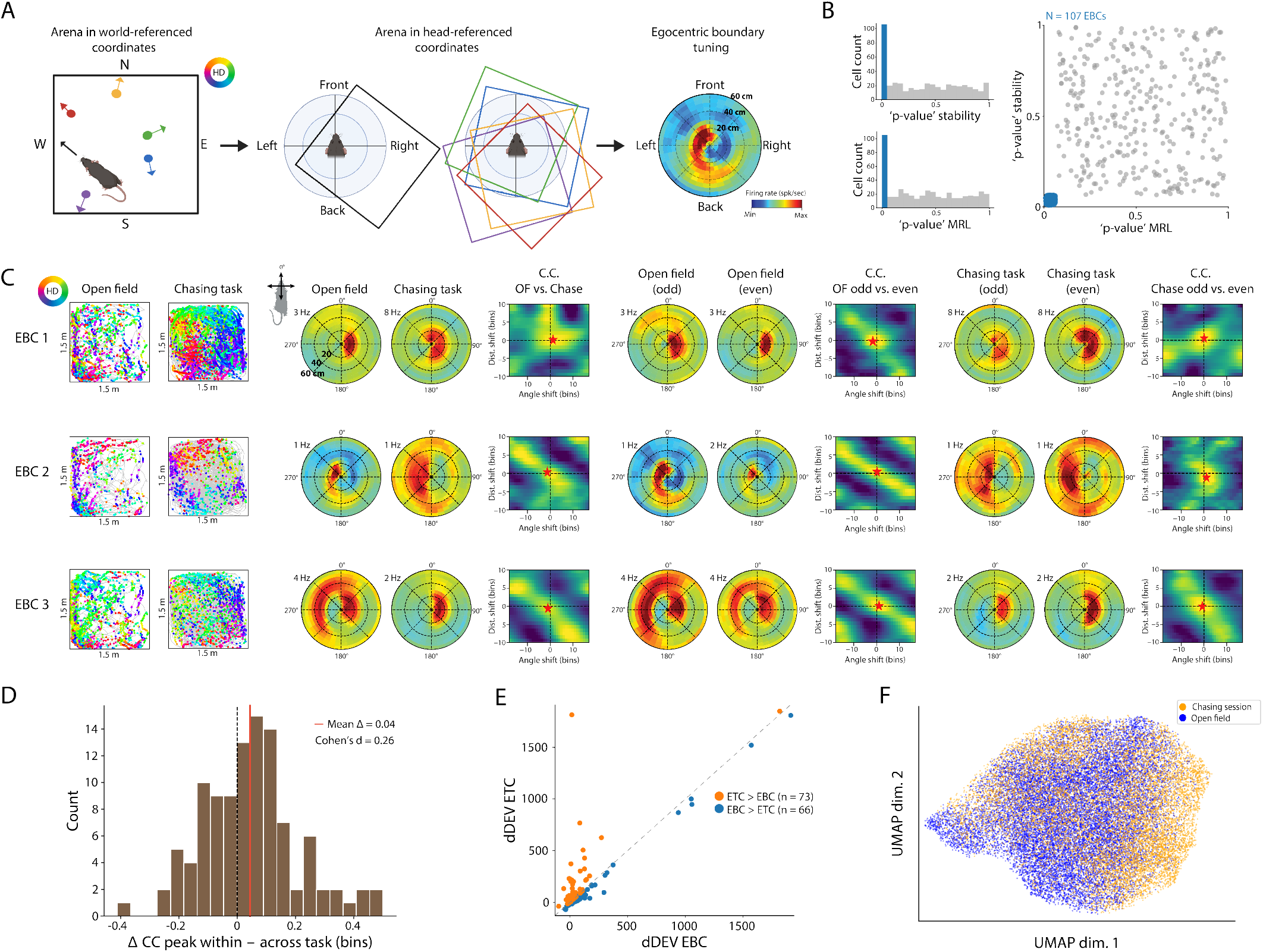
Egocentric boundary cells remain stable during pursuit. (A) Schematic for generating egocentric boundary rate maps (EBRs). Left, recording arena in world-referenced (allocentric) coordinates with head direction indicated by color. Middle: same arena shown in head-referenced (egocentric) coordinates, illustrating animal-referenced boundary positions from different locations in the arena. Right, example egocentric boundary rate map (EBR) showing firing rate as a function of the angle and distance (20–60 cm rings) of the arena wall relative to the head. (B) Selection criteria for EBCs. Upper Left, histogram of pseudo-*p* values for within-session stability. Lower Left, histogram of pseudo-*p* values for mean resultant length (MRL); blue bars indicate cells exceeding the 99th percentile of shuffled data for each measure. Right, joint distribution of MRL and stability pseudo-*p* values. Blue points (lower left corner) indicate neurons classified as EBCs. (C) Examples of egocentric boundary cells (EBCs). Each row for each neuron shows (left to right): the path of the animal with spikes overlaid; egocentric boundary rate maps (EBRs) for open field and chasing sessions; odd and even minute EBRs from each task; cross-correlograms (CCs) between odd and even minute EBRs. Cross correlogram peaks (red stars) shifted minimally both between- and within-task. (D) Distribution of cross-correlogram (CC) peak shifts for EBC–EBC cell pairs, expressed as the absolute shift (in bins) of the maximum cross-correlation peak. Blue histogram shows within–open field stability (OF even–odd), and orange histogram shows across-context comparison (OF vs chasing). Vertical dashed lines indicate the median peak shift for each distribution. Peak-shift distributions did not differ significantly between conditions (n.s.). (E) Scatter plot of deviance explained (dDEV) for GLM classifications as an EBC or ETC. Orange points correspond to neurons with larger dDEV for ETC than EBC models (*n* = 73), and blue points are neurons with greater dDEV for EBC than ETC models (*n* = 66). 28 cells were classified as both and 292 were neither; the diagonal illustrates the segregation between ETCs and EBCs (STAR Methods). (F) UMAP embedding of EBC population activity. Blue dots show open field time points; orange dots show chasing. In contrast to ETCs, which segregated in UMAP space, EBC population activity was more intermingled across behavioral contexts, indicating more stable population dynamics.

Based on these criteria, 107 of 459 neurons (23.3%) were classified as EBCs, identified first in the open field then verified for stability in the pursuit task. EBC tuning to boundary angles and distances remained highly consistent across behavioral tasks, though the firing fields could vary qualitatively in shape or size (Figure 2C). Analysis of cross-correlograms (CCs), however, confirmed the spatial stability of EBCs across tasks: peak shifts across chasing and open field tasks (0.45 bins) were slightly smaller than within-task drift (0.49 bins), reflecting a small effect size (Cohen’s d = 0.26) (Figure 2D).

To determine if switching tasks altered temporal structuring in the EBC network, we examined pairwise temporal relationships across 1,446 EBC–EBC cell pairs during open field and pursuit sessions. Temporal cross-correlations of binned spike trains revealed that the across-task change in peak correlation strength was near zero (median Δ = −0.004; Wilcoxon signed-rank test, *p* = 0.18)(Figure S4). Furthermore, pairwise coupling strength was strongly preserved across behavioral contexts (Spearman *ρ* = 0.44, *p* = 3.2 *×* 10^−10^). Finally, we compared the temporal lag of the cross-correlation peaks within open field sessions (even-odd split) and across tasks. The peak-lag varied less between tasks than within the even-odd comparison (∼ 1.3 s for open field vs. chasing; ∼ 1.8 s for open field even-odd split). On the whole, these findings provided no evidence of measurable spatial or temporal reorganization of EBC activity during active pursuit.

We further tested whether boundary and target encoding represent separable explanatory dimensions within individual neurons by quantifying the deviance explained (dDEV) by generalized linear models (GLMs) [38] fit separately to each variable (Figure 2E; STAR Methods). Rather than co-clustering, neurons dissociated into distinct groups either with boundary- or target-specific explanatory power, suggesting independent tuning dimensions rather than a single, flexible boundary code. Finally, used UMAP analysis to visually resolve whether chasing behavior impacted EBC population structure. In contrast to the strong reconfiguration seen in ETCs, time points from open field and chasing sessions did not form separate clusters (Silhouette score = 0.055; Dunn index = 0.093), indicating stable EBC population dynamics (Figure 2F). This stability was accompanied by higher overlap in state space across tasks (39.4% for EBCs vs. 12.1% for ETCs), and markedly lower LDA classification accuracy for behavioral context (66.2% for EBCs vs. 91.3% for ETCs).

Together, these analyses suggest that egocentric boundary representations in the RSC form a stable spatial scaffold that is maintained across different behavioral contexts. The preservation of pairwise temporal correlations and the invariance of EBC population dynamics in state space further support that pursuit-related ETC signals do not emerge from a retuning of the underlying boundary code.

### Egocentric coding of static objects

To determine if ETCs were specialized for moving targets or reflected generic coding of nearby objects [39, 40], we recorded RSC activity during sessions where a static object (10 *×* 50 cm striped cylinder) was placed in the southeast quadrant of the arena (Figure 3A). We tested for the expression of egocentric object cells (EOCs) using the same polar coordinate framework and thresholds applied to ETCs (MI and stability both above the 99th percentile of shuffled data). In the subset of recordings collected in object sessions (*N* = 3 rats, 402 neurons), 44 units (10.9%) were classified as EOCs (Figure 3B), which stably encoded the position of the stationary object relative to the head (Figure 3C). Notably, the spatial preferences of EOCs for objects had no relationship to the target location during pursuit (Figure 3C, rightmost column). This distinction was corroborated at the population level as well-UMAP embeddings provided visual evidence that EOC activity occupied different states during object sessions compared to chasing sessions (Figure 3D). This separation did not generalize to EBCs, which remained largely invariant across task contexts (18.1% overlap for EOCs v. 41.2% overlap for EBCs; Figure 3D). The EOCs’ lack of generalization between the object and the target (quantified in the following section) indicates that EOCs and ETCs are part of functionally segregated networks for coding stationary landmarks versus moving targets, respectively.

**Figure 3:**
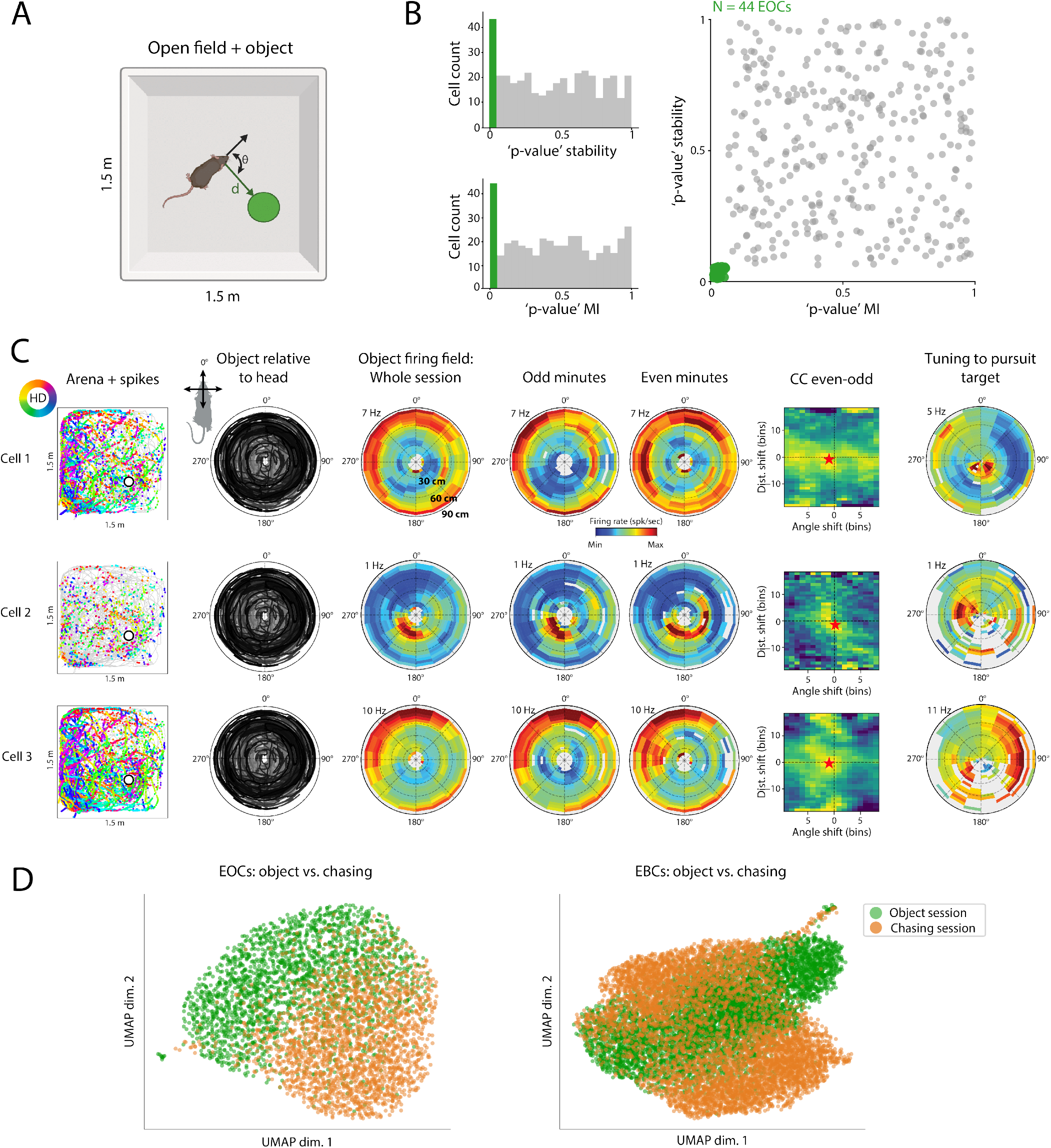
Egocentric object cells encode the location of a static object relative to the head. (A) Schematic showing the open field with static object in it. The egocentric position of the object is computed as the angle *θ* and distance *d* relative to the animal’s head direction. (B) Classification statistics of egocentric object cells (EOCs). Left, histograms of pseudo-*p* values for within-session stability (top) and spatial mutual information (MI, bottom), with orange bars indicating neurons exceeding the 99th percentile of their respective shuffled distributions. Right, joint distribution of MI and stability pseudo-*p* values; orange points indicate neurons classified as EOCs (*p* ¡ 0.01 for both criteria). (C) Example egocentric object cells (EOCs). Each row shows, from left to right, allocentric trajectory with spike positions colored by head direction and the object located at the open circle; egocentric object trajectory around the animal; egocentric coding of object location averaged for the whole session (rings indicate 30–90 cm distances); odd and even minute rate maps demonstrating stable tuning; cross-correlogram for odd and even minute rate maps (red stars denote CC peak); rate map showing the response of the same cell relative to the moving target during the chasing task (rightmost column). The firing patterns relative to the object do not carry over to the moving target during chasing. (D) Left, UMAP embedding of EOC population activity. Neural population states during the object session (green dots) segregated largely from population activity during the chasing task (orange dots) (Silhouette score = 0.23; Dunn index = 0.13; LDA accuracy = 81.8%). Right, EBC population activity from the same object and chasing sessions was substantially less separated (Silhouette score = 0.06; Dunn index = 0.096; LDA accuracy = 58.6%), indicating that EOCs distinguished the object and chasing conditions whereas EBCs did not.

### ETCs, EBCs and EOCs are three distinct egocentric populations

With three egocentric populations identified, we asked whether they were intermixed anatomically or organized topographically within the RSC. We analyzed the pool of neurons (*N* = 402) for which all three tuning classifications were applied, but histological reconstructions revealed that EBCs, ETCs, and EOCs were thoroughly intermingled at all depths along the recording probes. All three groups of cells spanned dysgranular and granular subdivisions, and there was no evidence of a dorsoventral topography relative to the surface (Figure 4A)[41], suggesting that the distinct coding properties of the cells were not attributed to regional differences in anatomical inputs.

**Figure 4:**
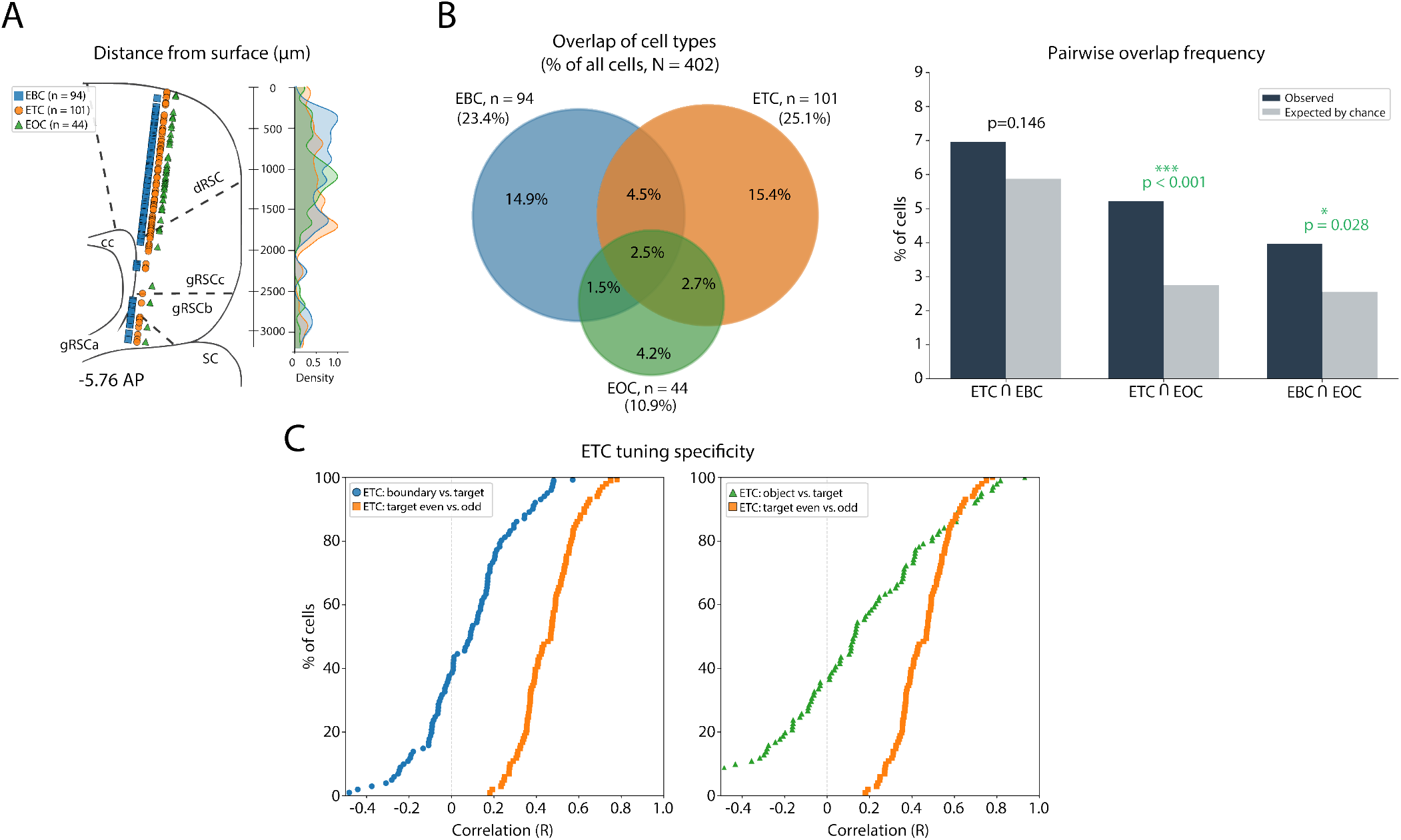
Three distinct but overlapping egocentric populations in RSC. (A) Egocentric cell types were distributed homogeneously in the RSC. Left, schematic coronal section showing reconstructed recording depths of each cell type relative to the cortical surface, summed across animals and overlaid. The schematic does not represent medial-lateral probe placement; functional cell types were grouped in their own columns for illustration purposes only (i.e. they are not different recording probes; EBCs shown as blue squares, ETCs as orange circles, EOCs as green triangles). Recording sites spanned dysgranular RSC (dRSC) and granular RSC subdivisions (gRSCa, gRSCb, gRSCc) and the corpus callosum (cc; in three rats). Passage through the cc led to a drop in cell yield from ca. 2,000–2,600 microns deep. Right, normalized density distributions of each cell type. Detailed histology, probe tracks and regional cell counts are shown per animal in Figure S1. (SC, superior colliculus). (B) Egocentric cell types consisted mainly of separate groups of cells. Left, Venn diagram showing the proportions of neurons classified as EBCs, ETCs, and EOCs. Percentages indicate both exclusive and overlapping classifications. Right, pairwise overlap frequency compared to overlap expected by chance. (C) ETC coding of target locations does not generalize to boundaries or objects. Left, cumulative distributions of spatial correlation coefficients for ETCs (orange squares) when rate maps for the target were measured against themselves during chasing vs. for arena boundaries (blue circles). Right, ETC tuning to the target was also significantly better correlated for within-session even-odd splits than for the target vs. object location.

The extensive intermingling of ETCs, EBCs and EOCs raised the possibility that the same cells multiplexed different egocentric variables. Although earlier UMAP visualizations indicated differences in population structure, such embeddings distort distances and cluster relationships, so are not definitive. A more direct test was whether individual neurons exhibited one or more egocentric codes. Overlap analysis revealed a largely modular organization (Figure 4B). Of the 402 neurons (101 ETCs, 25.1%; 94 EBCs,v23.4%; 44 EOCs, 10.9%), only a small minority exhibited multi-category tuning: 4.5% were classified as both EBC and ETC, 2.7% as both ETC and EOC, 1.5% as both EBC and EOC, and 2.5% met the criteria of all three (Figure 4B). Hypergeometric tests (Figure 4B, right) confirmed that the overlap between EBCs and ETCs did not differ significantly from chance (*p* = 0.146), further indicating their mutual functional independence. In contrast, the overlap of EOCs with ETCs and EBCs was modest but significantly greater chance (EBC–EOC: *p* = 0.028; ETC–EOC: *p <* 0.001), suggesting that static object representations overlap more at the neural level with boundary and target codes than those codes do with each other.

The functional segregation of EBCs and ETCs prompted us to test whether target and boundary information could both be extracted simultaneously from RSC population activity. Using a Bayesian decoder, we were able to reconstruct the animal’s egocentric bearing and distance to both the moving target and arena boundaries from RSC ensembles during pursuit (Video S3, Figure S5). Bait decoding was highly accurate, with median absolute distance errors of 0.096 m (mean = 0.123 m) and 18°(mean = 29.6°) for bearing, and both measures significantly exceeding chance levels (distance: *p <* 0.002, *z* = 3.14; bearing: *p <* 0.001, *z* = 5.63). Boundary decoding was similarly precise, with median errors of 0.133 m (mean = 0.18 m) for distance and 12.6°(mean = 18.8°) for bearing (distance: *z* = 15.21; bearing: *z* = 20.24, both *p <* 0.001). These successful reconstructions demonstrate that the RSC concurrently maintains parallel egocentric representations of both static boundaries and dynamic targets during active pursuit.

Given that target and boundary information could be decoded simultaneously from the same ensembles, we sought to verify whether the spatial tuning of ETCs was specific to the pursuit target and not other environmental features. We tested this by comparing the stability of target-centric rate maps of ETCs (even vs. odd segment split) against rate maps from the same cells referenced to the arena boundaries or static object. This revealed a clear divergence in coding: whereas ETC tuning was largely stable for the moving target (mean *r* = 0.46), target-centric rate maps showed a near-zero spatial correlation with arena boundaries (mean *r* = 0.06; *D* = 0.77, *p <* 0.001) and the static object (mean *r* = 0.14; *D* = 0.56, *p <* 0.001) (Figure 4C). Thus, ETC activity was selective for the moving goal. To determine if the target-centric coding required active engagement in chasing, we introduced the bait at an unattainable height above the arena (*>* 50*cm*) while mimicking the dynamics of a chasing trial. Strikingly, this passive observation failed to evoke the characteristic firing fields seen during typical chasing sessions (*N* = 44 ETCs from one animal; *D* = 0.55, *p <* 0.001; Figure S6), providing direct evidence that ETCs were not merely driven by the sight of the moving bait, but gated by active engagement in pursuit.

### RSC dynamically reweights behavioral variables during pursuit

Although ETC population activity was reorganized during pursuit, it remained unclear what this reflected mechanistically. We did not know, for example, whether the separate population states in UMAP were driven by global changes in excitability or a shift in the tuning properties of the neurons.

To address this, we classified neurons as excited, suppressed or indifferent during chasing (STAR Methods). Spike trains were binned at 50 ms resolution and firing rates were normalized by each neuron’s whole-session mean. We used a Welch’s *t*-statistic and circular shuffle testing (1000 iterations, preserving chasing bout number and duration) to compare normalized firing rates from chasing intervals against open field sessions. This identified subsets of chasing-excited and suppressed cells that rapidly swapped states at pursuit onset and offset in all four animals (Figure 5A), which was not likely explained by locomotor speed alone since the rate difference between between excited and suppressed cells was *>*2x larger during pursuit than high-speed epochs in the open field (Figure S7). Across the total population (*N* = 459), 69.9% of neurons were chasing-excited, 11.1% were suppressed, and 19% were indifferent (Figure 5B). These proportions– which were analogous within the ETC population specifically (Figure S8)– demonstrated that population state-shifts during pursuit were not due to global up- or down-modulation of spiking.

**Figure 5:**
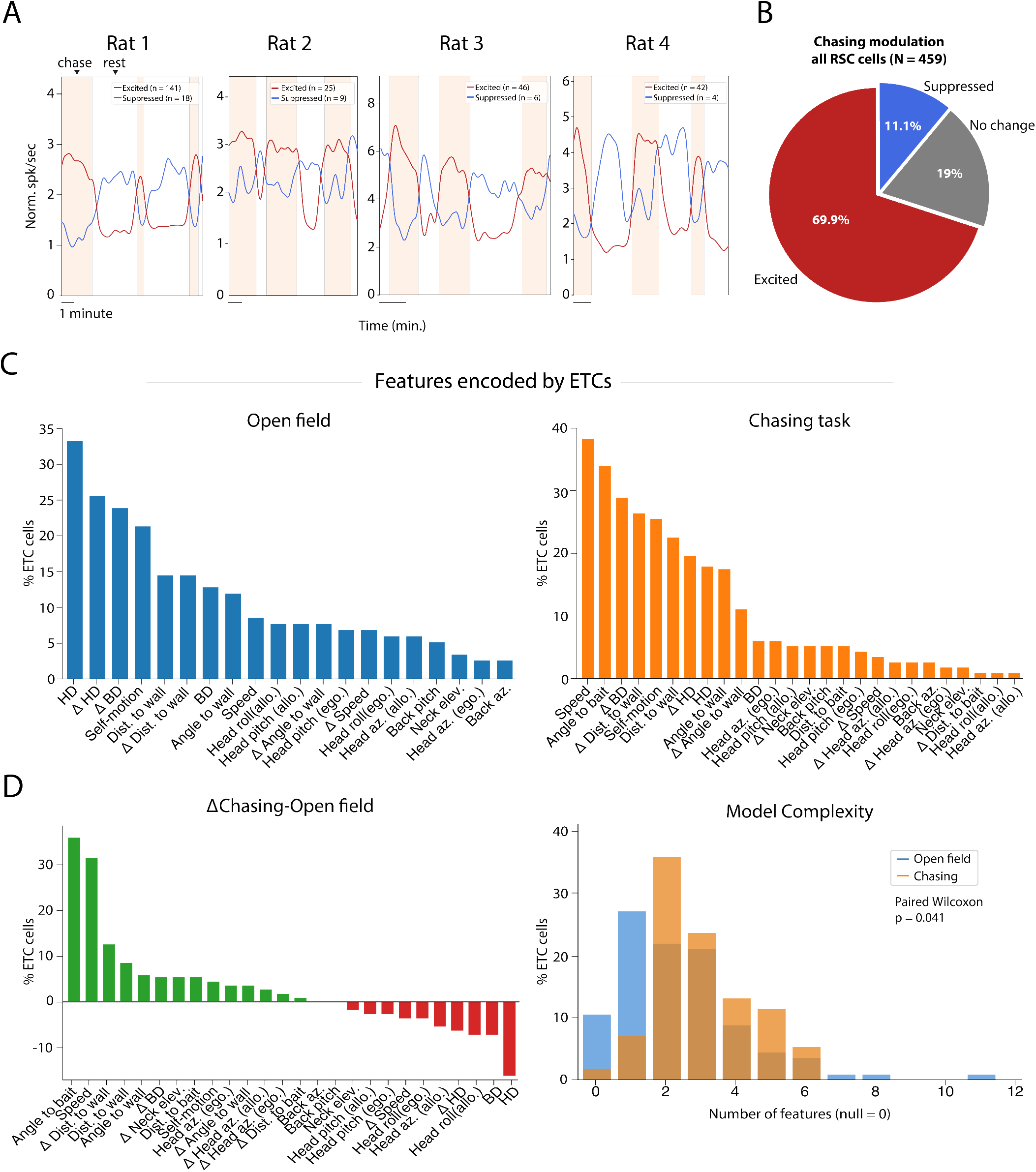
RSC dynamically reweights behavioral variables during pursuit. (A) Chasing-excited and suppressed cells switched states sharply at the onset and offset of chasing vs. consumatory rest periods. Each graph shows the session-normalized mean firing rate for chasing-excited (red line) and chasing-inhibited (blue) neurons (Methods) in each of four rats, with chase intervals indicated in shading and rest intervals in white. (B) Pie chart summarizing the fractions of chase response types across the total population of RSC cells. Nearly 70% of all RSC neurons were significantly up-modulated during chasing bouts, 11% were suppressed, and 19% were indifferent. (C) Bar graph showing the frequency with which specific behavioral features were included in the best-fitting GLMs for the ETC cell population (*N* = 101). Total counts exceed 100% because the best-fit models typically included more than one covariate. Left, in the open field, the most frequently selected features explaining ETC activity were allocentric head direction, changes in allocentric head/body direction, self-motion and egocentric bearing to the nearest wall. Right, same, but for chasing sessions, showing increased prevalence for running speed, bearing to the nearest wall, self-motion covariates and prominent encoding of the bait bearing relative to the head; allocentric head and body direction both decreased relative to the open field. (D) Left, the change in covariates selected by the GLM between contexts (chasing minus open field). Green bars indicate covariates included more frequently during chasing, red bars indicate covariates included less frequently. The two most prominently added features were relative angle of the bait and running speed, and the two features with the largest losses were allocentric head and body direction. Right, the distribution of best-model complexity *k* (number of covariates) for ETCs was significantly lower in the open field (blue) than in the chasing task (orange) (paired Wilcoxon signed-rank test: *p* = 4.14 *×* 10^−2^, *n* = 101).

The rapid changes in firing activity suggested strong modulatory effects of pursuit behavior, prompting us to assess whether chasing influenced *which behavioral variables* the cells encoded. We assessed this using GLMs [38] with forward-search model selection, which enabled data-driven identification of each neuron’s relevant covariates without the computational burden of an exhaustive combinatorial search [42]. This approach allowed us to compare which specific features best explained ETC spiking activity when animals transitioned between open field exploration and active chasing.

The GLMs included a comprehensive set of behavioral covariates (Figure 5C) including allocentric head/body direction and their derivatives, egocentric bearing and distance to the target or the walls, self-motion, running speed, turning velocity, as well as postural features and their derivatives (Methods)[43, 42]. ETC activity in both tasks reflected a mixture of ego- and allocentric variables, but allocentric head direction was the most commonly selected in the open field (Figure 5C left). During chasing, however, there was a robust increase in cells encoding speed, egocentric bearing to the target, and self-motion variables (Figure 5C right). A tally of the net differences in features selected across tasks (Figure 5D, left) highlighted a marked shift away from allocentric head and body direction in the open field toward self-referential and motoric features during chasing, especially egocentric bearing to the target and running speed, egocentric bearing to the walls and postural derivatives.

This shift in tuning properties was associated with an increase in the complexity of tuning structure, with ETC firing reflecting a richer combination of behavioral features when animals pursued the moving target, even with bait-angle and -distance subtracted from the covariate count (Figure 5D, right). Importantly, the reweighting of tuning properties and increased model complexity persisted when compared against high running-speed epochs in the open field (Figure S9), speaking against modulation solely by locomotor speed.

Together, these results demonstrate that RSC does not merely amplify egocentric signals during pursuit via gain modulation. Instead, ETCs undergo a dynamic reallocation of representational bandwidth, shifting away from allocentric features to egocentric variables critical for immediate action, providing a further basis for the population-level shifts in state space.

## DISCUSSION

The present study investigated how the retrosplenial cortex guides navigation when animals must simultaneously track a dynamic goal and the layout of the local environment. By recording neural populations during both open field exploration and naturalistic bait-chasing, we found that the RSC accomplishes this using a modular, parallel egocentric organization rather than a single overarching coordinate system. Whereas egocentric boundary cells stably encoded local boundary geometry across behaviors, egocentric target representations were recruited flexibly and re-tuned according to the demands of pursuit. This functional organization, where non-overlapping channels track boundaries and targets, points to the RSC as an active manager of representational resources rather than a passive coordinate translator. This enables the RSC to both maintain a stable scaffolding anchored to the environment and re-optimize dynamically for immediate action without mutual interference.

### Modular coding of more than open space

The use of open, static recording environments has been critical to establishing the foundations of allocentric spatial mapping in hippocampal–entorhinal circuits [44, 45, 46, 47], and identifying the likely role of the RSC in translating between allocentric and egocentric reference frames. More recent work uncovered robust coding of egocentric boundaries, self-motion, head movement and visual flow in the RSC,[28, 7, 8, 18, 25], but again in the context of exploratory behavior or passive movement. The critical step in the present work was additionally engaging the animals in chasing a dynamic goal, and explicitly comparing neural codes during exploratory tasks versus active pursuit. This revealed that egocentric coding in the RSC is not monolithic but modular, comprised of separate channels that represent diverse features in parallel.

We identified three functionally distinct populations– egocentric target cells (ETCs), egocentric boundary cells (EBCs), egocentric objects (EOCs)– that speak to a layered framework in the RSC. In it, we posit that EBCs provide a base layer anchoring the animal to the persistent spatial structure of the environment. Target representations occupy a flexible layer on top of the environmental scaffolding, and are recruited dynamically to meet the demands of pursuit. Although these populations are anatomically intermingled, they differ in tuning properties as well as population dynamics and task dependence, suggesting that they are not implemented by a single, generic transformation mechanism. Instead, they speak to the RSC maintaining multiple levels of signaling: a fixed environmental base plus additional levels for items the animal may encounter in that environment.

### Boundaries as persistent infrastructure

A notable feature of our results was the near-total stability of egocentric boundary representations across behavioral contexts: EBCs maintained highly consistent tuning and pairwise temporal cross-correlations during open field exploration and active pursuit (Figure 2D)– despite the heightened locomotor and cognitive demands of chasing. In this respect, EBCs may serve a role in anchoring egocentric coordinates analogous to that of HD and grid cells, which generate maps of allocentric space regardless of the specific environment or behavioral state[44, 46].

This stability also informs ongoing debates about allocentric–egocentric interactions in the RSC. Whereas previous work emphasized tight coupling between HD signals and egocentric representations,[11, 15] our results show that egocentric boundary coding can remain stable even when allocentric variables are down-weighted during pursuit. This suggests that boundary coding may be computed locally within RSC or inherited from parietal or other inputs[48, 36], rather than depending primarily on allocentric maps.

### Target coding as flexible resource allocation

In contrast to the stable scaffolding of EBCs, ETCs were context-dependent, with the majority of cells excited beyond baseline during chasing bouts (Figures 5A, S8). More revealing than rate modulation was the dynamic reweighting of behavioral covariates explaining ETC firing, wherein allocentric head direction was most common during exploration, and speed and egocentric target bearing were prioritized during the chase (Figure 5D). This difference persisted when compared against high-speed epochs in the open field (Figure S9), which speaks against locomotor speed as the primary explanation and raises an intriguing possibility: that a subset of neurons appearing as HD cells reference distal landmarks in the open field, but dynamically re-anchor to track the moving bait during pursuit. In this view, ETCs are a dynamically re-tunable subclass of cells distinct from stable EBCs. While we cannot confirm this possibility with existing recordings, future experiments can test directly whether cells swap their preferred anchoring online as the bait is introduced.

Regardless of the underlying circuit mechanism, the allo-to-ego reweighting indicates that ETCs reflect a state-dependent representational regime shaped by behaviorally urgent demands. During pursuit, heading relative to the “prey” becomes computationally more pressing than heading relative to the world, driving a transition from global spatial mapping to an action-centered view. Our present results can be thus be thought of as a reallocation of ‘representational bandwidth’ in the RSC, not unlike computational state changes observed, for example, when zebrafish switch from exploration to exploitation during prey capture [49]. Because neural populations have finite capacity, encoding all spatial variables with equal fidelity at all times would be inefficient. By dynamically adjusting representational priority, the RSC flexibly supports shifting behavioral functions without requiring entirely separate populations for every task. This view is supported by the increased model complexity observed for ETCs during chasing (Figure 5D), where the same cells take on richer computational demands during target pursuit and interception.

### Population dynamics suggest distinct operating regimes

Further evidence for a parallel architecture emerged at the population level, where ETC activity exhibited task-dependent state-space shifts that were notably absent in the EBC population (Figure 1F). This separation was robust across the open field and chasing tasks for ETCs (LDA accuracy = 91.3%), whereas EBCs were less affected (LDA accuracy = 66.2%), speaking to different operating regimes in each population. This observation, along with the single-cell findings above, suggests that target tracking engages *specific subsets* of cells in the RSC in a distinct computational mode rather than driving a region-wide transformation in all cells. This mode—which reflects specific changes in activity patterns rather than global gain modulation—could arise from task-dependent gating of specific inputs or shifts in recurrent dynamics that prioritize goaldirected pursuit over static environmental mapping.

### A family of egocentric codes

The identification of egocentric object cells (EOCs) extends this modular architecture beyond pursuit, revealing a family of egocentric codes tuned to distinct environmental elements. While EOCs stably represented a static landmark, they failed to generalize to moving targets (Figures 3C, D), reinforcing the functional segregation between stable features and dynamic goals. Overlap analysis (Figure 4B) showed that EOCs intersected significantly with both the spatially stable EBCs (*p* = 0.028) and dynamically-targeting ETCs (*p <* 0.001), whereas EBCs and ETCs were themselves independent (*p* = 0.146). This pattern suggests that EOCs may occupy a representational “middle ground” within the RSC. Whereas the EBC-EOC overlap could reflect that boundaries and objects both provide static reference points, the stronger overlap between EOCs and ETCs may suggest that discrete entities, whether static or moving, evokes common tracking computations. In this framework, object-related coding can interface with either stable boundary maps or dynamically gated target representations, enabling pursuit without disrupting global spatial scaffolding.

### Implications for navigation and goal-directed behavior

Together, these results prompt a reconceptualization of RSC function. Rather than acting solely as a coordinate transformation hub, the RSC appears to manage multiple egocentric representations and shift between predominantly allo- and egocentric modes of coding. This permits the maintenance of stable environmental scaffolding while allocating additional resources for behaviorally urgent goals. Such a flexible, modular organization suggests the RSC functions as a dynamic integrator of stable contextual signaling and flexible tracking of immediate goals.

### Limitations of the study

Several practical limitations have left several questions that remain open after this study. First, due to limitations in the number of targets we could physically deploy in the pursuit task, it is unclear whether ETCs comprise a single population of cells dedicated to tracking moving targets, or whether different targets would recruit additional, distinct subsets of ETCs. Second, our analyses focused on two-dimensional egocentric space, and it is not unlikely that extending the analysis to three dimensional space would reveal additional organizational principles [50, 51, 52]. Third, the bait-chasing paradigm, while ethologically motivated, involved experimenter-controlled bait movement rather than autonomous prey behavior. Real predatory pursuit involves prey that respond to the predator, which introduces action-reaction dynamics that were absent in our task. Future work using more naturalistic prey-like stimuli could address whether the representations we observed generalize to autonomous targets. Fourth, all animals in this study were female. Sex differences in spatial cognition have been documented in rodents [53], so for now it remains unknown whether the ETCs and EOCs described here differ between sexes. Finally, the circuit mechanisms underlying the stability of EBCs and the flexibility of ETCs remain unknown. Determining whether these properties arise from intrinsic RSC dynamics, upstream inputs, or top-down modulation will require causal manipulations. Such experiments will be essential for establishing how parallel egocentric coding contributes to navigation and decision-making.

## RESOURCE AVAILABILITY

### Lead contact

Requests for further information and resources should be directed to and will be fulfilled by the lead contact, Jonathan R. Whitlock.

### Materials availability

This study did not generate new unique reagents.

### Data and code availability

- The data collected in this study will be available from the corresponding author upon reasonable request.
- The custom written code developed for and used in this study will be made available on GitHub upon publication.
- Any additional information required to reanalyze the data reported in this paper is available from the lead contact upon request.

## ACKNOWLEDGMENTS

The authors warmly thank M. Andresen for critical contributions during surgeries, experiments, histological processing.; K. Haugen, K. Jøran Jenssen, and L. Hillisch for technical support; M. P. Witter and G.M. Olsen for helpful discussions and anatomical guidance; S. Eggen for veterinary oversight; members of the Whitlock lab for helpful discussion. Figure panels 1A, 2A and 3A, Created in BioRender, Saldanha, P. (2026) https://BioRender.com/kd8fkzs. This work was supported by the Research Council of Norway Investigator grants (No. 300709 and No. 759033), and through its Centres of Excellence scheme (project No. 332640, Centre for Algorithms in the Cortex), by National Infrastructure grants from the Research Council of Norway (NORBRAIN, projects number 295721 and 350201) and the Kavli Foundation.

## AUTHOR CONTRIBUTIONS

Conceptualization, P.S., J.R.W. and B.A.D.; methodology, P.S., M.B. and B.A.D.; investigation, P.S.; formal analysis, P.S. and M.B.; writing—original draft, P.S. and M.B.; writing—review & editing, P.S. and J.R.W.; visualization, P.S. and J.R.W.; supervision, J.R.W. and B.A.D.; funding acquisition, J.R.W.

## DECLARATION OF INTERESTS

The authors declare no competing interests.

## DECLARATION OF GENERATIVE AI AND AI-ASSISTED TECHNOLOGIES

During the preparation of this work the authors used Gemini v. 3.0 to edit and reduce word count in some sections of the original text. After using this tool, the authors reviewed and edited the content as needed and take full responsibility for the content of the published article.

## STAR Methods

### Experimental model and study participant details

#### Animals

Four adult Long Evans rats (female, 3-4 months of age) were used in this study (ID 28685-*Arwen*, 29865-*ToothMuch*, 29373-*PreciousGrape*, 30146-*MimosaPudica*). Animals were handled daily for ¿1 week prior to experiments. After handling, each animals was habituated to a black 140 *×* 140 *×* 50 cm recording arena for several days before surgery and recordings. The recording environment consisted of the recording box, a white cue card on the north wall, and recording hardware setup beyond the south wall. The recording environment remained consistent across all experiments in the study. The animals were screened for chasing behavior prior to experiments, so were already familiar with the task during recordings. The behavioral conditions consisted of open field foraging in the empty arena, in which the animal foraged for corn puff crumbs sprinkled on the arena floor by the experimenter. Chasing sessions were in the same area, and included a fishing rod with a food reward (a corn puff, either bare or smeared with Oreo cream or Nutella) suspended on the end of a string, and moved pseudorandomly by the experimenter to evoke chasing. A retroreflective spherical marker was attached 9 cm above the bait for tracking during chasing trials. The object session was the same as open field foraging. Recording sessions were 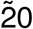 minutes, and chasing sessions were repeated until achieving ¿20 minutes of active chasing behavior for three animals and 7 minutes in the fourth (#28685).

All experiments were performed in accordance with the Norwegian Animal Welfare Act, the European Convention for the Protection of Vertebrate Animals used for Experimental and Other Scientific Purposes, and the local veterinary authority at the Norwegian University of Science and Technology. All experiments were approved by the Norwegian Food Safety Authority (Mattilsynet; protocol ID 25094). The study contained no randomization to experimental treatments and no blinding. Sample size (number of animals) was set a priori to at least four, given the expected cell yield necessary to perform unbiased statistical analyses.

### Method details

#### Surgery

Animals were anesthetized initially with 5% Isofluorane vapor mixed with oxygen and maintained at 1-3% throughout the surgical procedure. When anesthetized, they were placed in a stereotaxic frame and injected subcutaneously with Metacam (2.5mg/kg) and Temgesic (0.05mg/kg as general analgesics. Body temperature was maintained at 37°C with a heating pad. A local anesthetic (Marcain 0.5%) was injected under the scalp before the first surgical incision was made, followed by clearing the skull of skin and fascia. Following the clearing of tissue, adjustments were made to make sure the skull was leveled before a high-speed dental drill was used to open a craniotomy over the retrosplenial cortex (Kulzer GmbH, Germany). A single jeweler’s screw was inserted over the cerebellum as a ground and reference screw for the probe implant, connected with a silver wire and sealed with SCP03B (silver conductive paint). Animals were subcutaneously administered fluids at the start, at the half-way point of surgery and immediately post-surgery.

A single-shanked silicon probe (Neuropixels version 1.0, IMEC, Belgium) was stereotaxically lowered in the right hemisphere of two animals and left hemisphere for other two animals at coordinates AP: -5.1-6.0, ML: 0.7-0.9, DV: 4.9-6.0. The craniotomy and unimplanted length of the probe were enclosed with a silicone elastomer (Kwik-Sil, World Precision Instruments Inc., USA). After allowing the silicone elastomer to harden, a thin layer of Super-Bond (Sun Medical Co., Ltd., Japan) was applied to it and the skull to increase bonding strength between the skull surface and dental acrylic. To minimize light induced electrical interference during the experiments the first layer of dental cement was dyed black and care was taken to completely cover any light-sensitive parts of the probe. Undyed cement was further applied to secure the probe. A custom 3D printed skull cap was used to encase the implant outside the brain, serving both as a protective housing and as a base for attaching the rigid body for tracking the head in 3D during recordings. The cap was secured to the skull with black-dyed cement. Once the cement had cured, sharp edges were removed with the dental drill at the end of surgery. Following surgery, the animals were placed in a heated (32°C) chamber to recover. Once awake and ambulating, they were returned to their home cage and allowed food and rest before data collection commenced.

### Recording procedures

Electrophysiological recordings were performed using Neuropixels 1.0 acquisition hardware: A National Instruments PXIe-1071 chassis and PXI-6133 I/O module for recording analog and digital inputs. Implanted probes were operationally connected via a headstage circuit board and interface cable above the head. An elastic string was used to counterbalance excess cable, allowing animals to move freely through the entire recording area during recordings. Data acquisition was performed using SpikeGLX software (SpikeGLX, Janelia Research Campus), with the amplifier gain for AP channels set to 500x, LFP channels set to 250x, an external reference and AP filter cut at 300 Hz. Spike activity was recorded and collected from bank 0, i.e., the most distal 384 recording sites. Behavioral experiments and neural recordings were performed 532 a minimum of 2.5 hours after animals had fully recovered from surgery.

### 3D tracking and synchronization

Animals had circular marker cutouts of 3M retro-reflective tape (OptiTrack, catalog no. MSC 546 1040; Natural Point Inc., Corvallis, OR, USA) placed on their body at the neck, back and tailbase, which were shaved during surgery. To capture head rotations we used a rigid body with four retro-reflective spheres, size 7.9 mm, 9.5 mm and 12.7 mm (OptiTrack, catalog no. MKR079M3-10, MKR095M3-10 550 and MKR127M3-10, respectively) integrated with the skull cap. The markers were tracked retroreflectively using 8 near-infrared (NIR) recording cameras (7 NIR, and one black/white camera for referencing; Flex 13 cameras, Part #: FL13, OptiTrack, Oregon, USA) at 120 FPS in optical motion capture software (Motive version 2.2.0; OptiTrack).

Tracking and neural data streams were syncrhonized using three LEDs affixed to the sides of the recording arena, and captured by the motion capture system. The LEDs were controlled by an Arduino Microcontroller with C++ code to generate irregularly timed digital pulses (250 ms duration of each flash; random 250 ms ≤ IPI ≤ 1.5 sec) transmitted to acquisition systems throughout the recordings. Individual markers were labeled using built-in functions in Motive on raw .tak files. After recordings were completed, 3D data was reconstructed by creating a marker set for the three trunk markers, a rigid body construction for the capture of the animal’s head and a rigid body was constructed to capture the synchronizing LEDs. Using Motive’s built-in “autolabel” function, markers were identified and labeled, with remaining unlabeled markers manually labeled by the experimenter. Errors in marker assignment were corrected manually. For frames where markers were occluded, tracked points were reconstructed using in-built functions in the Motive software. Different modes of reconstruction (“cubic”) were used depending on the duration of occlusion, the behavior of the animal, marker identity and the potential for validating the reconstruction. After reconstruction and labeling of the markers, the positional information from each session was exported as a .csv-file. A custom-written Python script was used to process and align tracking information with the synchronization stream into a .pkl file. This file was then loaded in a custom graphical user interface (GUI) to construct a head coordinate system and merge the tracking data with spiketimes into a final .mat file. These same procedures are described in further detail in ([42]). The minimum threshold for determining the animals’ running speed was 5 cm/s.

### Perfusion and probe placement verification

Upon completion of the recordings, each rat was euthanized with an overdose of Isoflurane. This was followed by intracardial perfusion using saline and 4% paraformaldehyde. The probe shank was kept inside the brain tissue and the entire skull was kept in 4% paraformaldehyde for 24 hours. The next day, the brain was then carefully extracted from the skull and transferred to a solution of 2% dimethyl sulfoxide (DMSO, VWR, USA) for cryoprotection for a period of one to two days before cryosectioning. Following cryoprotection, the brain was frozen and sectioned coronally into three series of 40 *µ*m slices using a freezing sliding microtome (Microm HM-430, Thermo Scientific, Waltham, MA). The first series of sections was directly mounted onto Superfrost slides (Fisher Scientific, Gö teborg, Sweden) and subsequently stained with Cresyl Violet. All brain sections were digitized using a digital scanner and appropriate illumination wavelengths with scanning software provided by Carl Zeiss AS (Oslo, Norway). The resulting images were then visualized using ZEN (blue edition) software. Post-hoc probe reconstruction was done using HERBS[41].

### Chase interval annotation

Chase intervals were identified post-hoc during visual inspection of video recordings. An interval was classified as “chase” when the animal actively reoriented and initiated movement toward the bait, as indicated by locomotion direction, head orientation and sustained forward movement. After the animal caught the bait, the rod was removed from the arena; between-chase rest periods consisted of the animal consuming the food reward, as well as stationary grooming, rearing and undirected exploration. Two experimenters independently annotated and agreed on chase interval boundaries.

### Egocentric rate map construction

Egocentric rate maps were constructed by computing the position of spatial features (arena boundaries, bait, or stationary objects) relative to the position and direction of the animal’s head at each time bin. Spike data were binned at 8.33 ms (∼ 120 Hz). For each time bin, egocentric angle *θ* was computed as the angular offset between the feature and the animal’s head direction, and distance *d* was computed as the Euclidean distance from the animal’s head to the feature. For egocentric boundary cells (EBCs), rays were cast at 1^◦^ intervals from −180^◦^ to 180^◦^ relative to head direction, and distance to the nearest wall was computed along each ray. These distances were then averaged into 10^◦^ bins, yielding 36 angular bins.

For all egocentric cell types, rate maps were computed by accumulating occupancy and spike counts into a polar grid. Angular bins spanned −180^◦^ to 180^◦^ in 10^◦^ increments (36 bins). Distance bins spanned 0–60 cm in 5 cm increments for EBCs (12 bins) and 0–90 cm in 4.5 cm increments for ETCs and EOCs (20 bins). Firing rate in each bin was computed as spike count divided by occupancy time. Rate maps were smoothed using a 2D Gaussian kernel (kernel size: 3 *×* 3 bins; standard deviation: 1.5 bins). Bins with fewer than 50 frames of occupancy (∼ 0.4 s) were excluded from analysis.

### Quantification and statistical analysis

#### Skaggs information index calculation

To test whether a particular neuron was tuned to a dynamic target or not, we used the Skaggs information index [35, 54]. The information index is an approximation of the mutual information metric and measures the amount of information the firing of one neuron carries relative to some observed variable. The mutual information between two variables *X* and *Y*, I(*X*; *Y*), is given as

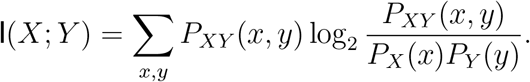

We let *X* denote the event that the animal is currently at a distance *d* from the target, with an angle *θ* between the target and the animal’s current heading direction, and let *Y* be the event that a neuron fires exactly *f* spikes at the current time. Considering binned spike data, the approximation then allows the information index to be written as

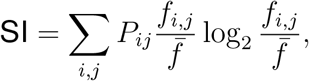

where *P*_*i,j*_ denotes the probability of being in the *i,j*-th binned distance-angle bin, *f*_*i,j*_ denotes the calculated firing rate in the *i,j*-th bin and 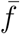 denotes the overall mean firing rate of the neuron.

#### Cell type classification using the Skaggs information index and stability

The Skaggs MI was selected for ETC and EOC classification because, unlike the angular-only MRL metric, it quantifies information across the full two-dimensional (distance–angle) egocentric manifold. Furthermore, because the bait was predominantly positioned directly ahead of the animal during pursuit, MI provided a more robust measure of tuning by explicitly weighting information content by the probability of occupancy in each spatial bin. For ETCs, egocentric distance–angle coordinates were defined relative to the moving target during chasing bouts; for EOCs, coordinates were defined relative to the stationary object during object sessions.

Tuning stability was quantified as the Pearson correlation coefficient between egocentric rate maps generated from alternating 30-s blocks (even-odd split) of the relevant behavioral epochs, ensuring that the spatial structure of the head-centered firing field was preserved throughout the recording session.

Significance thresholds for both measures were established for each neuron by circularly shifting its binary activity vector (1000 iterations) to generate cell-specific null distributions against which observed values were converted to pseudo-*p* values (0-1 scale). This normalization was necessary because baseline firing rates and temporal dynamics could vary substantially across neurons, causing the raw null distributions for MI to differ in scale for each cell (Figure S2); pooling raw scores resulted in a non-interpretable mixture of distributions. Converting scores to pseudo-*p* values ensured a standardized comparison across the recorded population: under the null hypothesis of no spatial tuning, pseudo-*p* values should be uniformly distributed. Significant spatial signals are thus represented by a non-uniform enrichment of neurons in the lowest *p*-value bins, exceeding the uniform density expected by chance (e.g., Figure 1D). For consistency, the same normalization was performed for Pearson’s correlations.

Neurons were formally classified as ETCs or EOCs only if both their Skaggs mutual information index and Pearson stability correlation exceeded the 99th percentile of their respective shuffled distributions (pseudo-*p <* 0.01). The shuffle and stability procedures were identical to those used for EBC classification (below), only with the mean resultant length (MRL) replacing the Skagg mutual information index as the primary tuning metric boundary-related activity.

#### Egocentric boundary cell classification using mean resultant length

The mean resultant length (MRL) was used as our tuning metric for classifying neurons as egocentric boundary cells (EBCs). The MRL measures the concentration of firing around a preferred egocentric angle and is computed as:

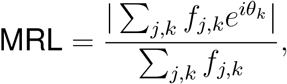

where *f*_*j,k*_ denotes the firing rate in the *j, k*-th distance-angle bin, *θ*_*k*_ is the *k*-th egocentric angular bin in radians, and *i* is the imaginary unit.

Statistical significance of the MRL was assessed using a shuffle procedure, where the spike train was circularly shifted by a random offset (minimum 30 s) and the MRL was recomputed. This was repeated 1000 times to generate a null distribution, and a neuron passed the tuning criterion if its observed MRL exceeded the 99th percentile of the null distribution (*p <* 0.01).

As with ETCs and EOCs, temporal stability was assessed by splitting the session into alternating 30-second blocks (odd and even) and computing separate rate maps for each half. The Pearson correlation between the odd and even rate maps was computed, excluding bins with ¡ 400 ms cumulative occupancy, and a shuffled null distribution was generated by circularly shifting spike trains and recomputing the odd even correlation (1000 shuffles; same as for ETCs and EOCs). A neuron passed the stability criterion if its observed correlation exceeded the 99th percentile of the null distribution (normalized pseudo-*p <* 0.01). Neurons were classified as EBCs if they passed both the MRL and stability criteria (sequential testing following Alexander et al.[7]).

#### GLM covariate analysis

We fit generalized linear models (GLMs) with a Bernoulli family and logit link function to predict binarized spike counts (spike vs. no spike per time bin) from behavioral covariates (as done in Mimica et al.[42]). Covariates included allocentric head direction and body direction and their first temporal derivatives, running speed and its derivative, angular velocity of body and head, egocentric bearing to target (relative head angle), egocentric distance to target (relative distance), posture variables[43, 42] including head azimuth, head roll, head pitch, back pitch, neck elevation, and their temporal derivatives.

For each neuron, we used a forward model selection[55] procedure to identify the best-fitting combination of covariates. Starting from a null (intercept-only) model, covariates were iteratively added one at a time. At each iteration, all remaining covariates were tested as candidate additions. Each candidate model was evaluated using 10-fold cross-validation, in which each fold contained temporally distributed sub-blocks to reduce autocorrelation effects. Model parameters were estimated using L-BFGS optimization with L1 regularization (*λ* = 0.0001, *α* = 1).

For each fold, we computed the log-likelihood ratio (LLR) between the candidate model and the null model, normalized by the number of spikes in the test set (LLR per spike). A candidate model was considered significantly better than the current best model if a one-sided Wilcoxon signed-rank test comparing cross-validated LLR-per-spike scores across folds yielded *p <* 0.01. Among all candidates meeting this significance threshold, the model with the highest mean LLR per spike was selected as the new best model. The forward selection procedure continued until no candidate model produced a significant improvement, at which point the current best model was returned.

This procedure was performed separately on the same cells for open field and chasing sessions to allow direct comparison of covariate importance across behavioral contexts. Model complexity *k* was defined as the number of covariates in the best-fitting model. Covariate usage frequencies and model complexities were compared between open field and chasing sessions using paired Wilcoxon signed-rank tests.

#### Deviance explained (dDEV)

To quantify the relative contribution of boundary-versus target-tuning for each neuron, we computed the deviance explained (dDEV) from GLMs. For each neuron, we fit separate GLMs predicting spike counts from (1) egocentric boundary position (angle and distance to nearest wall) and (2) egocentric target position (angle and distance to bait). Deviance explained was computed as:

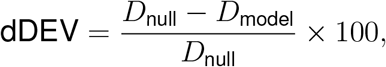

where *D*_null_ is the deviance of an intercept-only model and *D*_model_ is the deviance of the full model including spatial covariates.

#### Spatial cross-correlations between cell types

To quantify the similarity of the different types of egocentric tuning (relative to boundaries, chasing bait or object), we computed pairwise Pearson correlations between egocentric rate maps. For neurons classified as multiple cell types (e.g., both EBC and ETC), we correlated the EBC rate map (computed from open field data with respect to boundaries) with the ETC rate map (computed from chasing data with respect to the bait). Within-session stability was computed by generating correlations between rate maps for odd and even segments for a given session.

Correlations were computed only over bins with sufficient occupancy in both rate maps (minimum 50 overlapping bins). Rate maps were smoothed with a Gaussian filter (*σ* = 1 bin) before measuring correlations.

#### Temporal cross-correlations between cell pairs

To examine functional coupling between EBCs, we computed pairwise cross-correlations of binned spike trains. Spike trains were binned at 150 ms and detrended using a high-pass filter (window = 50 bins, polynomial order = 2) to remove slow drift. Cross-correlation coefficients were computed for lags spanning *±*60 bins (*±*9 s):

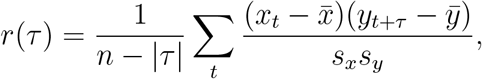

where *x* and *y* are the normalized spike trains, 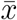 and 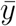 are their means, *s*_*x*_ and *s*_*y*_ are their standard deviations, and *τ* is the lag.

Cross-correlations were computed separately for open field and chasing sessions to assess whether functional coupling was preserved across behavioral states.

### UMAP projections

UMAP (Uniform Manifold Approximation and Projection)[37] was employed to visualize population activity in a low-dimensional space. First, binned spike trains were smoothed using a Gaussian filter with standard deviation *σ* = 3 bins and converted to firing rates (Hz). The smoothed spike trains were then downsampled to every 12th bin. Before UMAP embedding, firing rates were rank-normalized within each neuron using a Gaussian inverse transform, and principal component analysis (PCA) was applied to reduce dimensionality to 6 components.

UMAP was then applied with parameters: n neighbors = 100, min dist = 0.8, metric = ‘euclidean’. For comparing behavioral modulation in different tasks, UMAP was fit on concatenated data e.g. from both open field and chasing sessions, with time points colored by session type. Separate UMAP embeddings were computed for different cell populations (EBCs only, ETCs only) to assess how the different task conditions affected the state spaces occupied by each population.

### UMAP quantification

To quantify the degree of separation between behavioral contexts in UMAP embeddings, we computed several complementary metrics. The Silhouette score measures how similar each point is to its own cluster compared to other clusters, ranging from −1 (poor clustering) to +1 (perfect clustering):

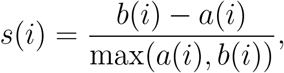

where *a*(*i*) is the mean distance from point *i* to other points in the same cluster and *b*(*i*) is the mean distance to points in the nearest other cluster.

The Dunn index (DI) measures overall clustering quality; a higher Dunn index corresponds to tighter clustering in the data, with 0 being no clustering. For it, centroids were first calculated for every behavioral cluster. Distances between each point within a behavioral cluster and the cluster’s centroid (intra-cluster distances) and the distances between centroids of different clusters (inter-cluster distances) were measured. The DI was then calculated as the ratio between the minimum inter-cluster distance and the maximum intra-cluster distance (as defined above). We computed a robust variant using the 5th percentile of inter-cluster distances to reduce sensitivity to outliers.

Context overlap was quantified as the percentage of time bins in the UMAP embedding that were closer to the centroid of the opposite behavioral context than to their own. Linear discriminant analysis (LDA) and support vector machine (SVM) classifiers were trained to predict behavioral context from UMAP coordinates using 5-fold stratified cross-validation.

### Cell type overlap analysis

To quantify overlap between cell type classifications, we computed the percentage of neurons meeting criteria for multiple cell types (e.g., both EBC and ETC). Statistical significance of overlap was assessed using hypergeometric tests, which compare the observed overlap to the expected overlap under the null hypothesis of independent classification:

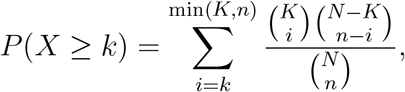

where *N* is the total number of neurons, *K* is the number of neurons in class A, *n* is the number of neurons in class B, and *k* is the observed overlap.

### 0.1 Bayesian decoding of egocentric wall and bait variables

To quantify the information carried by the RSC population about egocentric spatial variables, we implemented a Bayesian decoding framework to reconstruct independently the animal’s distance and bearing to (i) the nearest arena wall and (ii) the moving bait.

#### 0.1.1 Egocentric state variables

At each tracking frame (Δ*t* = median(Δ*frame times*), approximately 8.33 ms at 120 Hz), two egocentric coordinate pairs were computed. For the *bait encoder*, distance 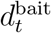 was the Euclidean distance from the animal to the tracked bait position, and bearing 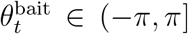 was the allocentric angle to the bait minus the animal’s head direction. For the *wall encoder*, we computed the distance 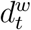 and egocentric bearing 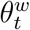 from the animal to each of the four arena walls (*w* ∈ {West, East, South, North}), using the perpendicular vector from the animal’s position to the wall boundary relative to head direction. Animal positions and head directions containing NaN values were cleaned by linear interpolation (positions) or unwrap-interpolate-rewrap (head direction) prior to covariate computation. Both distance and bearing signals were then temporally smoothed with a Gaussian kernel (*σ* = 100 ms / Δ*t* samples) using <monospace>scipy.ndimage.gaussian filte</monospace>

#### 0.1.2 Occupancy-normalized egocentric rate maps

For each neuron *i* and each target variable (wall or bait), we estimated a two-dimensional rate map 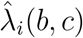 over a discretized (*d, θ*) grid. Spike times were converted to frame indices and only frames falling within the training mask and with finite covariate values were included. Occupancy Occ(*b, c*) was computed as the number of frames spent in each bin multiplied by Δ*t* (yielding seconds), and the spike histogram Spk_*i*_(*b, c*) was the corresponding spike count. The occupancy-normalized rate map was:

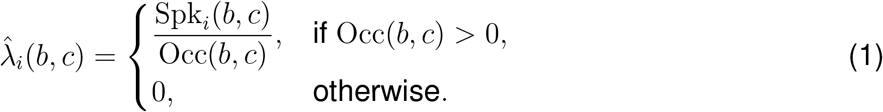

No additional spatial smoothing was applied to the rate maps, ensuring that decoding was driven by the raw spatial information of the population. For the wall encoder, distance bins spanned 0– 1.0 m in 0.1 m increments (*B*_*d*_ = 10 bins) and angular bins spanned the full 360^◦^ in 5^◦^ increments (*B*_*θ*_ = 72 bins). For the bait encoder, distance bins spanned 0–2.5 m in 0.15 m increments (*B*_*d*_ = 17 bins) with the same angular resolution (*B*_*θ*_ = 72 bins, 5^◦^ each).

#### 0.1.3 Poisson maximum-likelihood decoder

Decoding was performed in non-overlapping temporal windows of duration *W* = 0.2 s. Only windows in which all frames fell within the test mask and had finite covariate values were included. For each valid window starting at time *t*, we computed the spike count 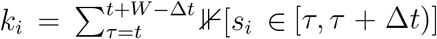 for each neuron *i*, where *s*_*i*_ denotes the set of spike times. Assuming conditional independence across neurons and Poisson-distributed spiking with expected rate 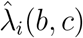, the log-likelihood for candidate bin (*b, c*) was:

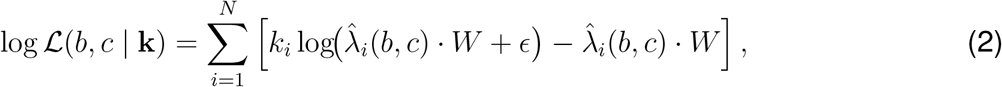

where *N* is the number of simultaneously recorded RSC neurons and *ϵ* = 10^−12^ prevents numerical underflow. The decoded position was taken as the maximum-likelihood estimate under a uniform prior over the (*d, θ*) grid:

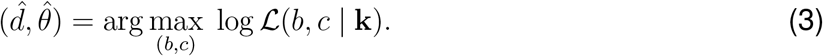

Decoded distance and bearing values were assigned at the bin centres of the winning bin.

#### 0.1.4 Train-test partitioning

For each animal, chase intervals from recording session 1 (c1) were used for training and chase intervals from recording session 2 (c2) were used for testing. Crucially, the wall and bait decoders were each fit estimating its own set of rate maps in its respective covariate space, but using the identical neural population and temporal partitions. This design ensures that any difference in decoding accuracy can be attributed solely to how much information the population carries about each spatial variable, rather than to differences in the neurons sampled or the data used for training and testing. Analyses were restricted to RSC neurons, identified by their recording channel position within histologically verified RSC boundaries.

#### 0.1.5 Accuracy metrics

Decoding accuracy was quantified using median absolute error and Pearson correlation. For distance:

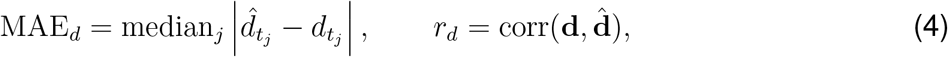

where the median is taken over all valid test windows *j*. For angular bearing:

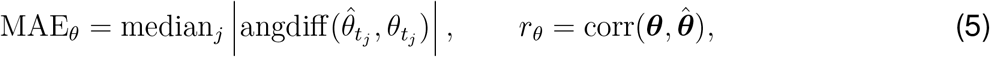

where angdiff(·, ·) denotes the signed circular difference wrapped to [−*π, π*].

#### 0.1.6 Statistical significance via circular spike-shift nulls

To assess decoding significance while preserving the temporal autocorrelation of individual spike trains, we generated null distributions by independently and circularly shifting each neuron’s spike train by a random amount drawn uniformly from [10 s, *T*_max_ −10 s], where *T*_max_ is the maximum spike time across all neurons. For each of *M* = 500 shuffles, rate maps were reestimated from the shifted spike trains and decoding was repeated on the test set. One-sided *p*-values were computed as the fraction of shuffled correlation values exceeding the observed correlation.

## Supplemental Figures and Legends

**Figure S1:**
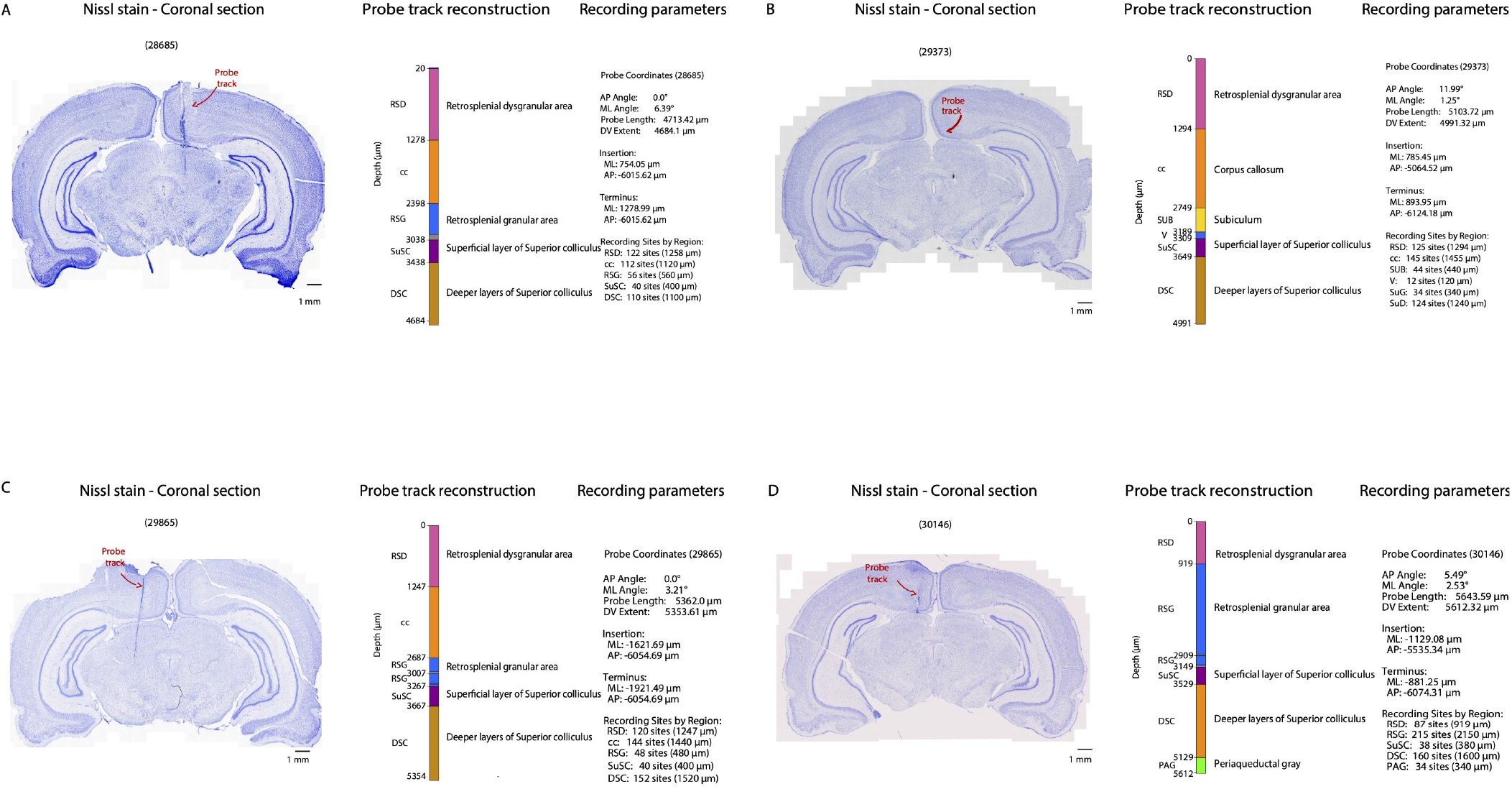
Histological verification of Neuropixels probe placements, related to Figure 1. (A–D) Probe track reconstructions for each of the four animals used in the study (IDs: 28685, 29373, 29865, 30146). For each animal, the left panel shows a Nissl-stained coronal section with the probe track indicated (red arrow). Center panels show the reconstructed probe trajectory from HERBS software[41], with recording sites assigned to cytoarchitecturally defined regions along the dorsoventral axis. Abbreviations are: dysgranular retrosplenial cortex (RSD), corpus callosum (cc), granular retrosplenial cortex (RSG), superficial layers of superior colliculus (SuSC), and deeper layers of superior colliculus (DSC). Additional regions were identified in individual animals, including subiculum (SUB) and periaqueductal gray (PAG). Right panels list stereotaxic coordinates, probe insertion and terminus positions, probe angle, and the number of recording sites per region. Depth (in *µ*m) and regional boundaries are marked along each reconstruction. Scale bars: 1 mm.

**Figure S2:**
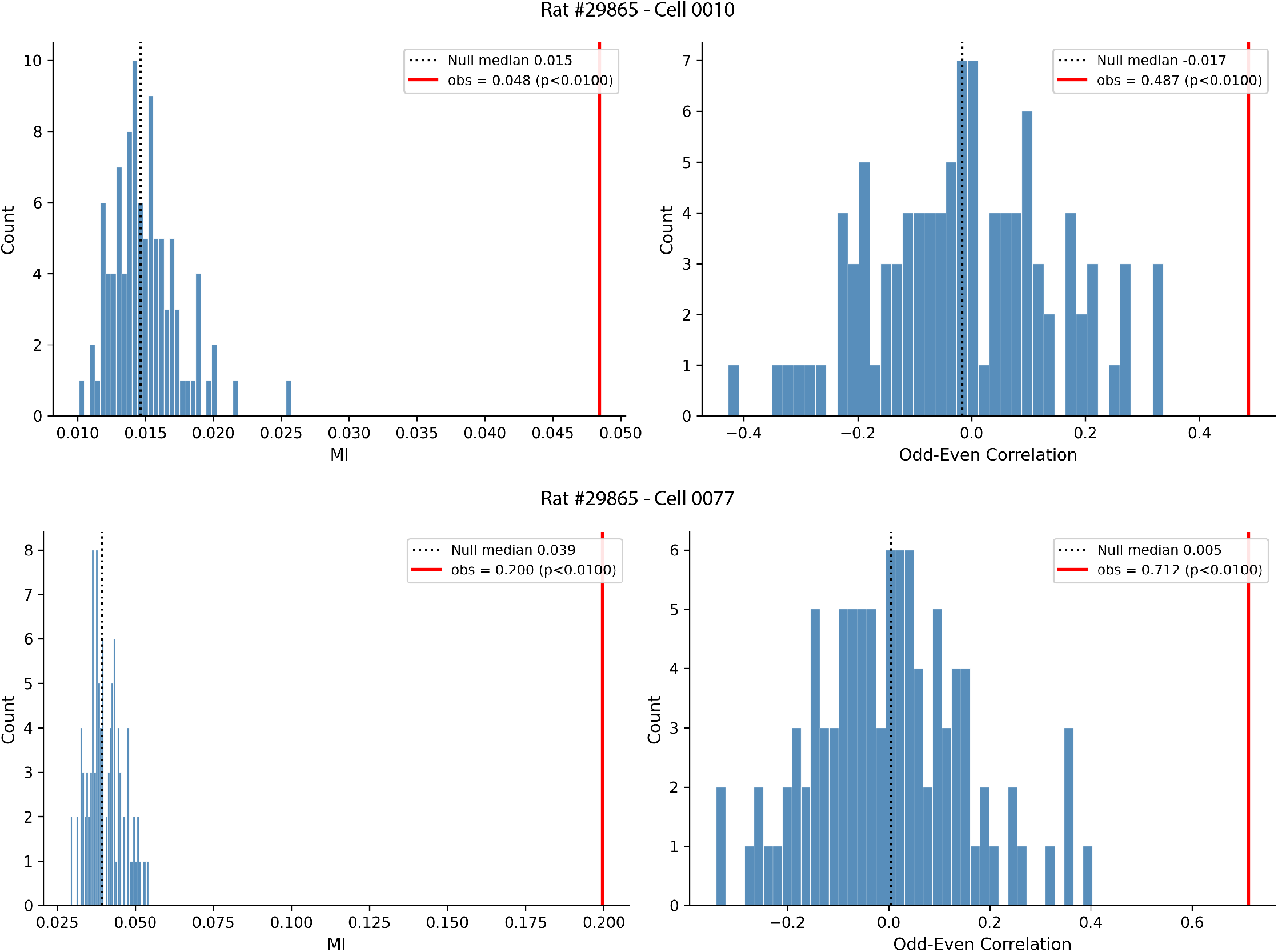
Examples of shuffled distributions for Skaggs MI and Pearson’s correlations for two example ETC neurons, related to Figure 1. (Top) Example of an ETC neuron whose Skaggs MI and stability measures (red line) both exceeded the 99th percentile of the shuffled distribution (blue bars). The observed Skaggs MI value for the top cell is far higher than its own shuffled distribution, but falls within chance levels for the second example cell (below). Because of the differences in scale, null distributions were expressed on a 0-1 scale per cell and converted to pseudo-p-values, with the significance threshold of *p <* 0.01.

**Figure S3:**
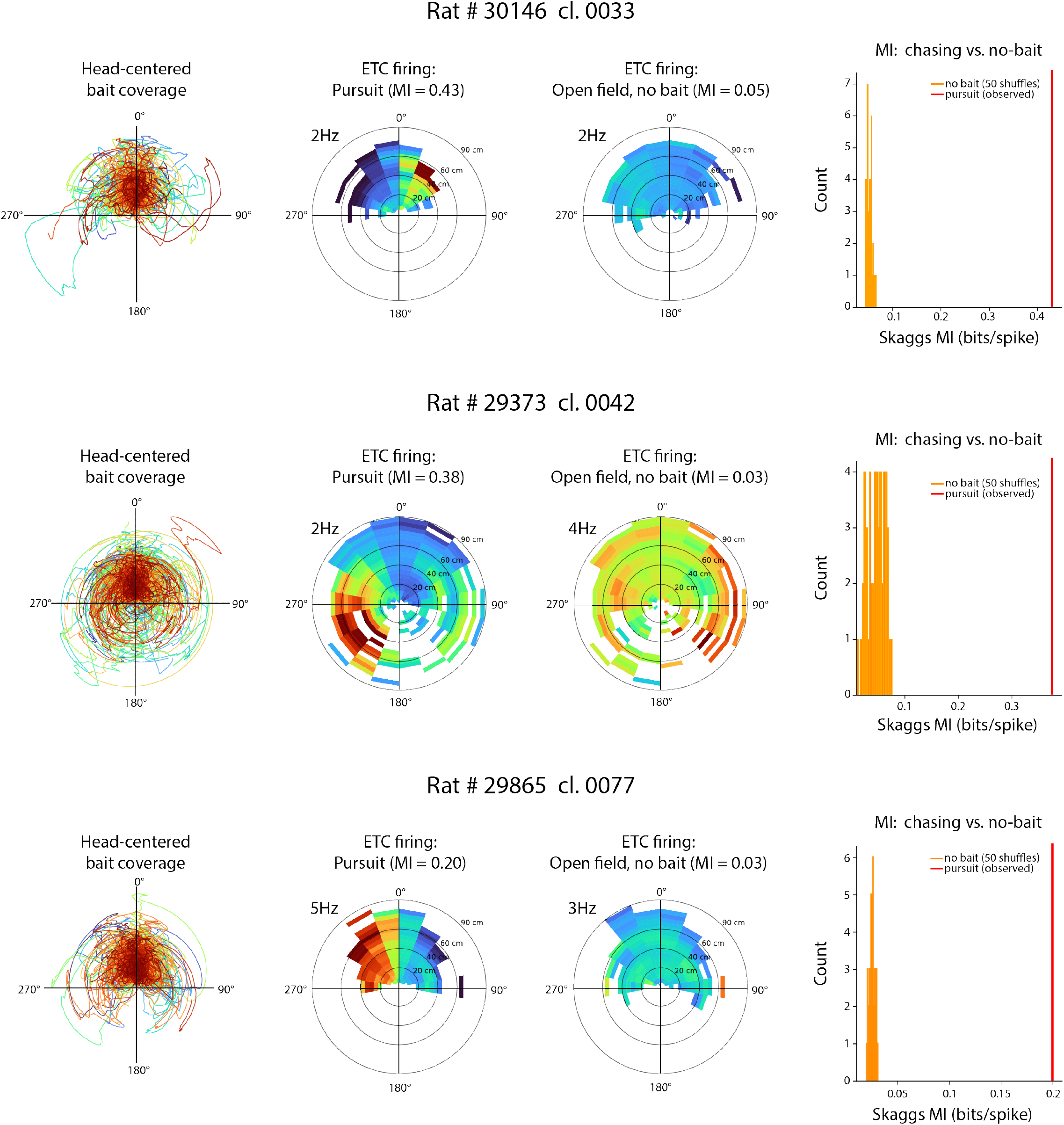
ETC firing fields are not expressed without a bait, related to Figure 1. Three examples demonstrating ETC firing fields expressed when the bait (left) was present during the pursuit task (middle left), but not not in the open field when the bait was absent (middle right). Rate maps for the open field used a “phantom” bait which occupied the same angle-distance bins sampled during the pursuit session, but assigned randomly to time bins from the open field. The rate maps were generated using open field spikes plus the “phantom” bait positions 50 times with random assignments taken from a real pursuit session.. Mutual information values were significantly higher (p¡0.001) for actual vs. “phantom bait” rate maps for all comparisons (right column).

**Figure S4:**
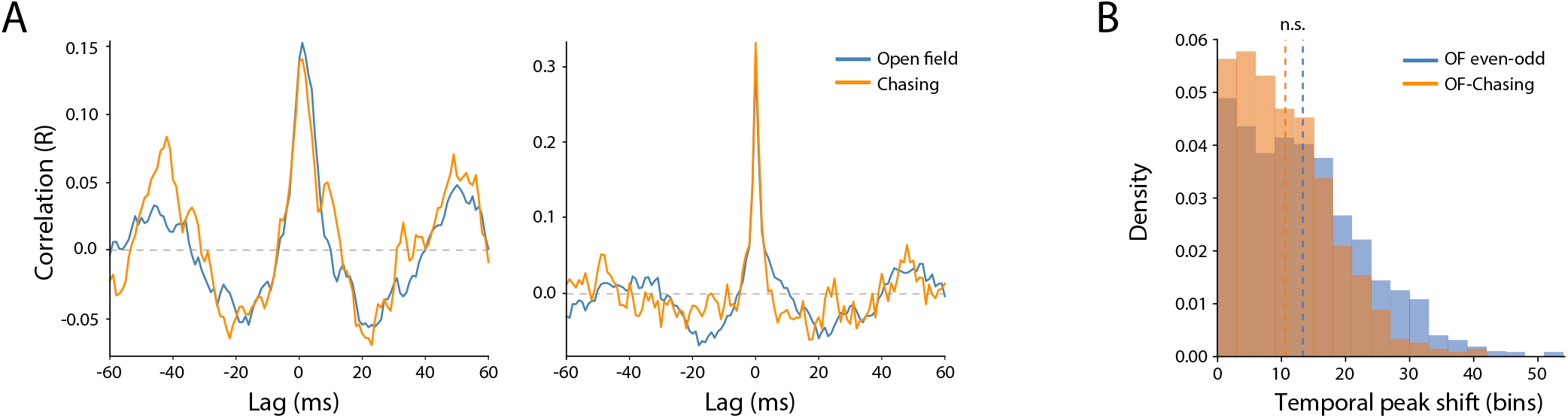
Temporal coupling between EBC pairs is preserved across open field and chasing behavior, related to Figure 2. (A) Example temporal cross-correlation functions for representative EBC–EBC cell pairs during open field exploration (blue) and chasing behavior (orange). Cross-correlations were computed from binned spike trains, which showed similar peak structure across behavioral contexts. (B) Distribution of temporal peak shifts across EBC–EBC pairs. Peak-lag differences were compared between open field sessions (even -odd split; blue) and across behavioral contexts (open field vs. chasing; orange). Peak-lag variability was comparable or lower across tasks (∼ 1.3 ms for open field vs. chasing) than within the open field session (∼ 1.8 ms for the even -odd split), indicating that changing tasks did not affect temporal coordination. Across 1,446 EBC–EBC pairs, the change in peak correlation strength between conditions was centered near zero, and pairwise coupling strength was strongly preserved across behavioral contexts (Spearman *ρ* = 0.44, *p* = 3.2 *×* 10^−10^).

**Figure S5:**
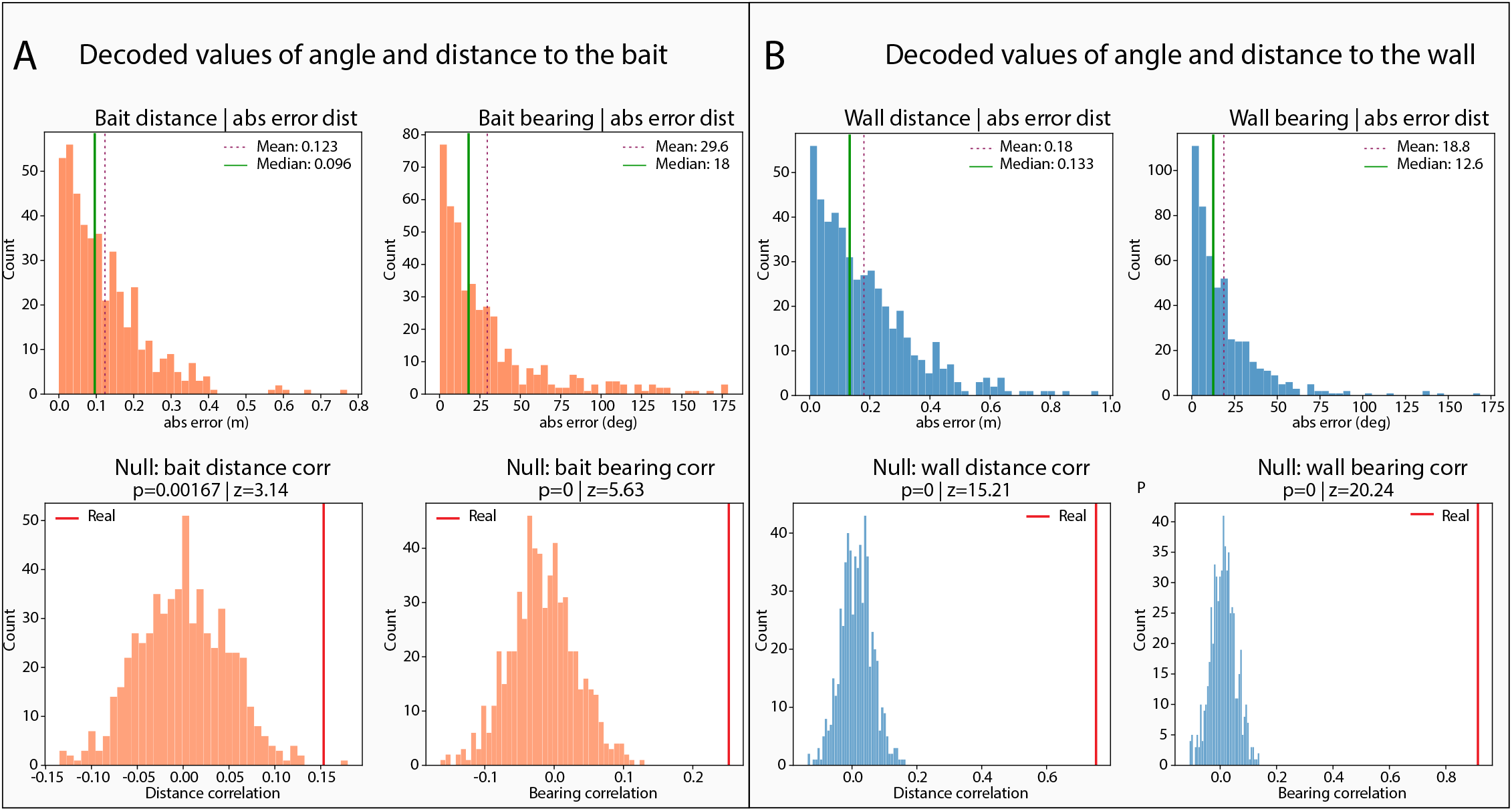
Simultaneous decoding of egocentric distance and bearing to walls and bait during chasing, related to Figure 4. (A) Decoding performance for egocentric distance and bearing to the moving bait during chasing sessions. Histograms show absolute decoding error for bait distance (left) and bait bearing (right). Mean and median errors are indicated by dashed and solid lines, respectively. Bottom panels show null distributions obtained by circularly shifting spike trains relative to behavior. Red vertical lines show actual decoding performance. Both bait distance and bearing correlations significantly exceeded chance levels. (B) Decoding performance for egocentric distance and bearing to the arena walls during chasing sessions. Histograms show absolute decoding error for wall distance (left) and wall bearing (right); mean and median values are indicated by dashed and solid lines, respectively. Bottom panels show null distributions for wall decoding. Real decoding correlations (red lines) were significantly greater than chance for both distance and bearing (both *p <* 0.001; *z* = 15.21 for distance and *z* = 20.24 for bearing). Together, this demonstrates that RSC population activity contains decodable information about both static environmental boundaries and dynamic targets in parallel during pursuit behavior.

**Figure S6:**
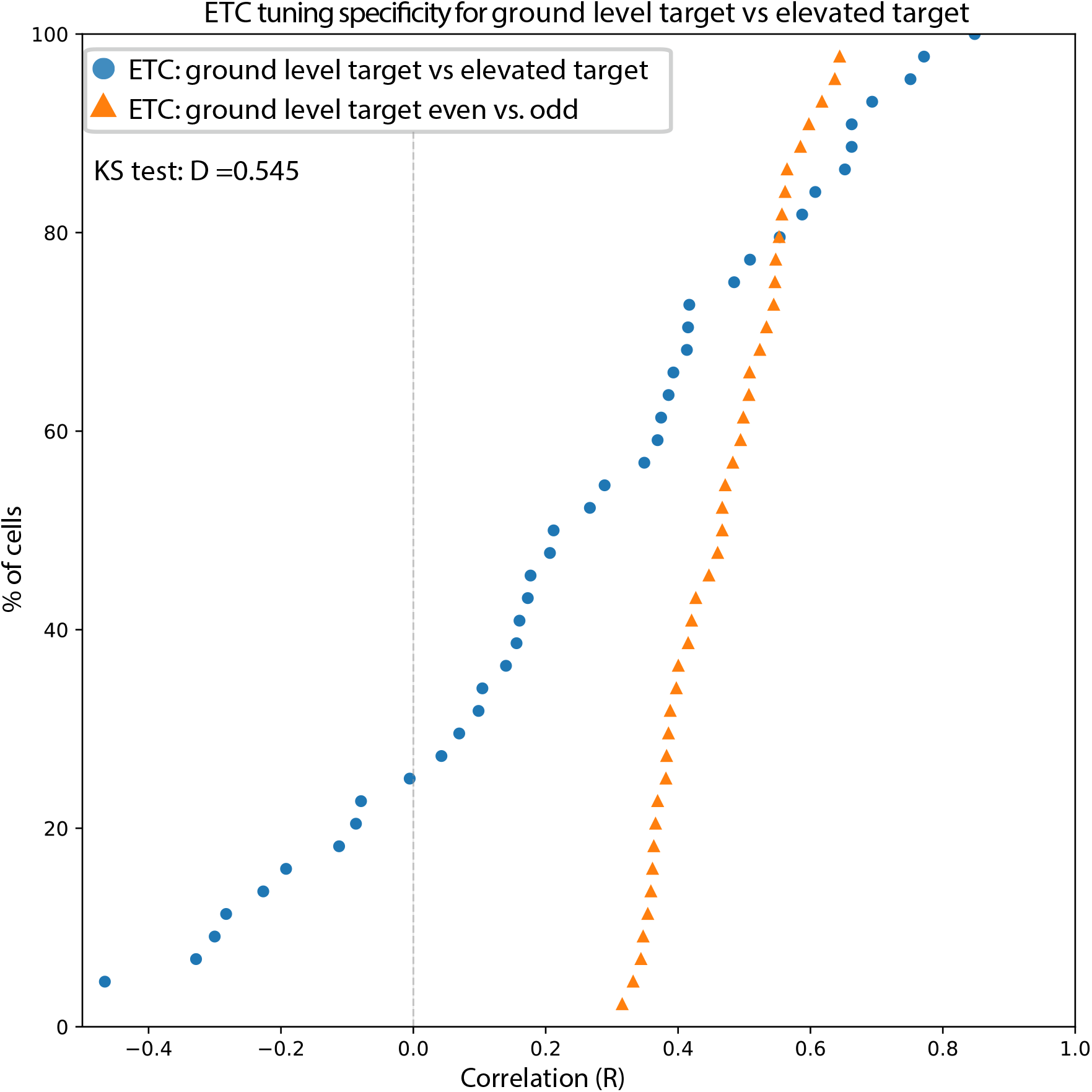
ETC spatial tuning is not evoked by visual presentation of the bait alone, related to Figure 4. Cumulative distributions of spatial firing-rate correlations for ETCs from one animal during active pursuit versus when the bait was above the arena. Correlations were computed between egocentric target-centered firing-rate maps in each condition. Orange triangles show within-session stability (even-odd split) during the pursuit session; blue circles show correlations between maps for ground-level pursuit against those with the target elevated. Spatial correlations were substantially higher for rate maps made with the bait at ground level than across bait elevation conditions (*R* = 0.48 for ETC even-odd vs. 0.23 for ETC-ETC’; Kolmogorov– Smirnov test: *D* = 0.545; *p <* 0.001), indicating that ETC tuning is specific to active engagement in pursuit of the ground-level target.

**Figure S7:**
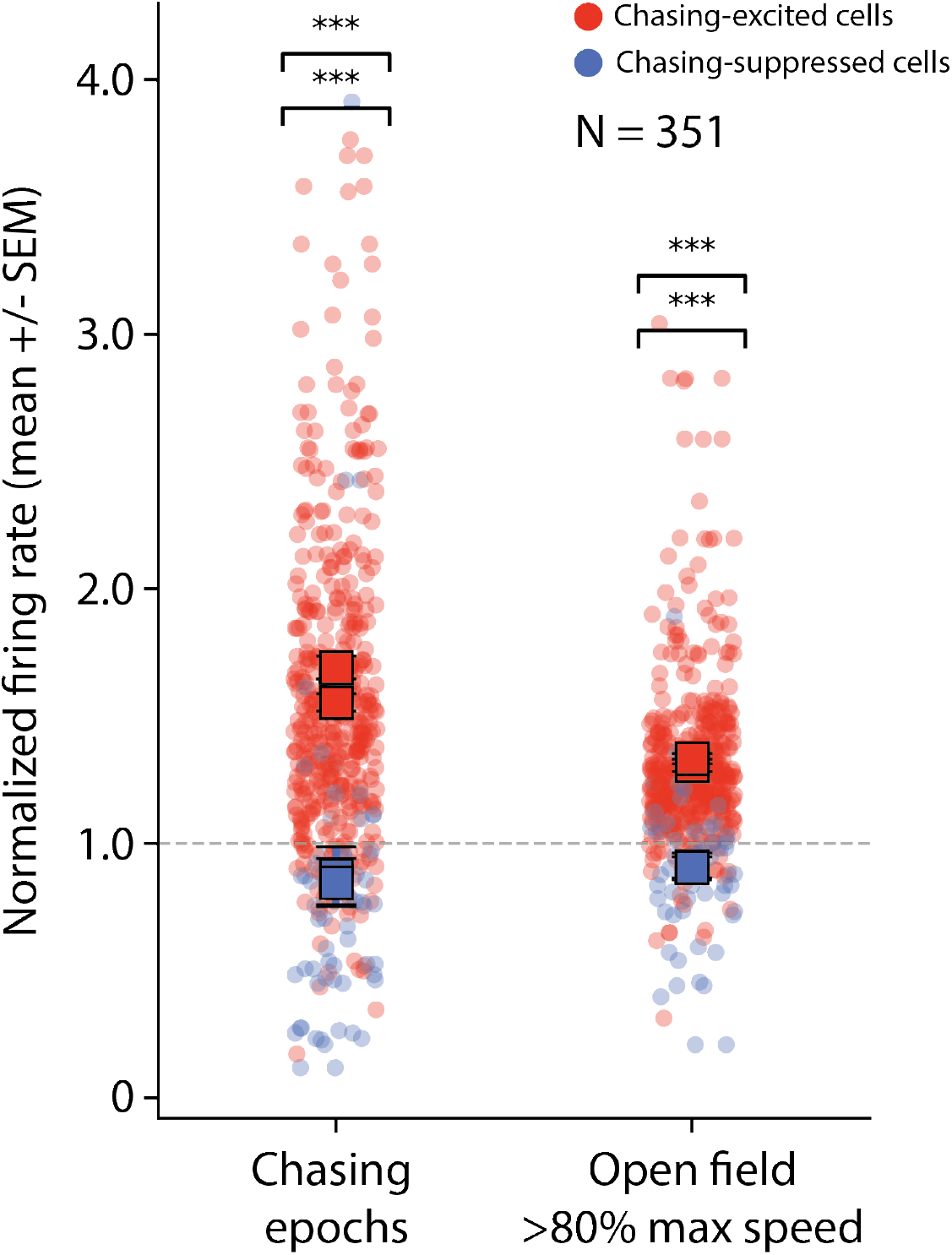
Firing rate modulation in the RSC is stronger during pursuit than high-speed epochs in the open field, related to Figure 5. Whisker plots showing baseline-normalized rate changes for the same chasing-excited and -suppressed cells (pooled across animals) during an example chasing session (left) and corresponding open field session (right). Open field data were taken from segments where animals exceeded 80% of the maximum speed within-session. Both excitation and suppression were stronger in the chasing task, with a *>* 2x larger difference between excited and suppressed cells during pursuit than in the open field. Box plots depict the median, first and third quartiles of each dataset.

**Figure S8:**
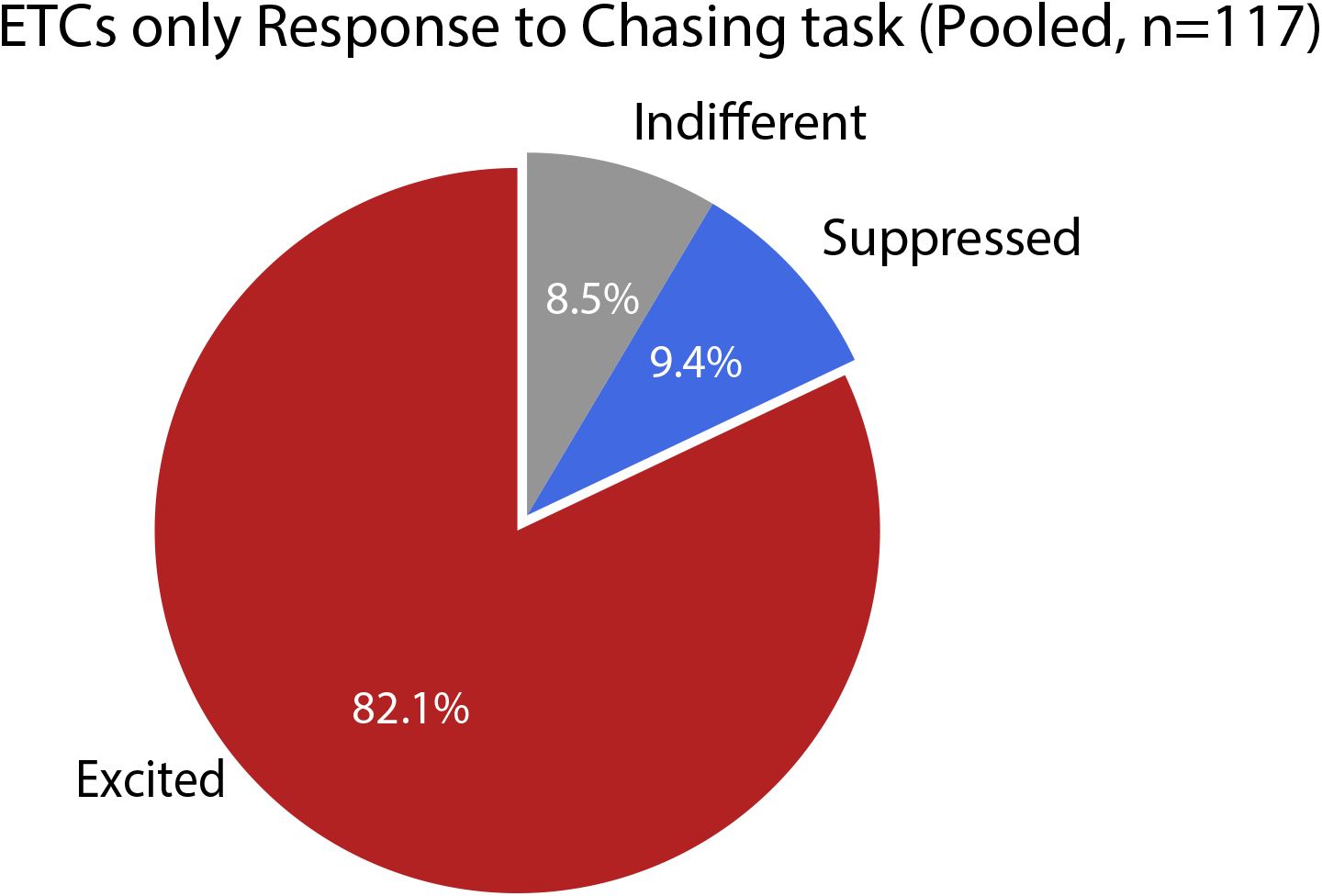
ETC rate modulation during chasing, related to Figure 5. Pie chart showing the distribution of excited, suppressed and unchanged ETCs during chasing behavior, pooled across sessions. Cells were classified based on changes in firing rate during chase intervals relative to non-chase periods. The majority of ETCs were significantly up-modulated during chasing, whereas smaller fractions were suppressed or showed no significant change.

**Figure S9:**
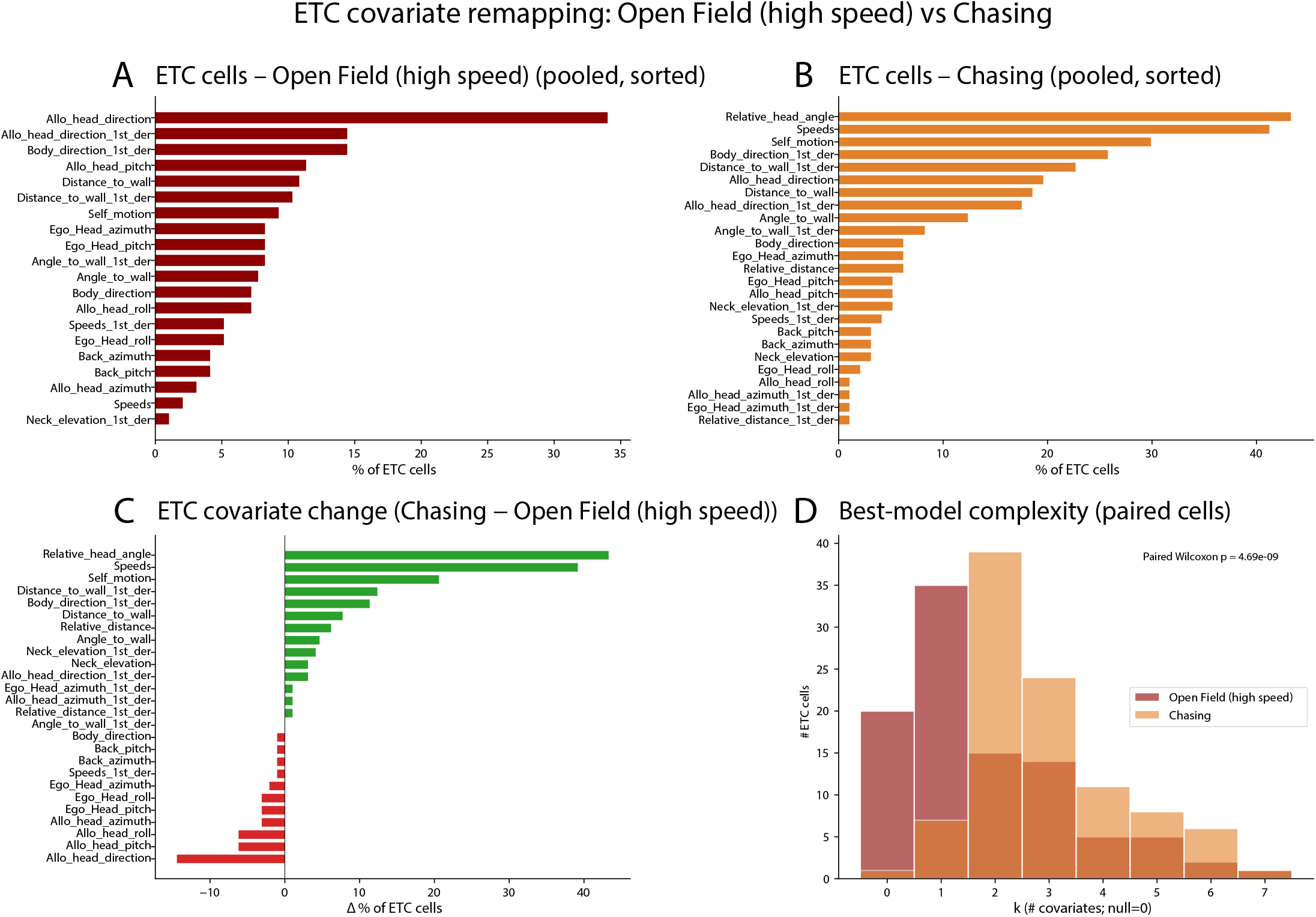
ETC covariate changes persist between chasing and higher-speed segments in the open field, related to Figure 5. (A) Distribution of covariates included in the best-fit models for ETCs during high-speed epochs in the open field, pooled across cells and sorted by prevalence. “High speed” epochs are those in which locomotor speed was in the upper 50% of running speeds measured in the open field (a 50% max cutoff was used to ensure adequate sampling for meaningful 10-fold cross validation). Bars show the percentage of ETCs whose best-fitting model included a given behavioral covariate. Allocentric head direction and body direction derivatives were among the most common features in open field. (B) Distribution of best-model covariates for the same ETC population during chasing behavior. Relative head angle, speed, and self-motion variables became more prevalent along with target-related covariates. (C) The net change in covariate prevalence between conditions (chasing minus high-speed open field) showed that target-bearing and other egocentric variables (speed, self-motion in particular) showed the largest increases (net increases shown as green bars), whereas head direction decreased the most (net losses shown as red bars), followed by world-referenced postural variables. (D) Distribution of best-fit model complexity (number of covariates) for ETCs paired across conditions (*N* = 79). Model complexity increased significantly during chasing compared to high-speed epochs in the open field (paired Wilcoxon test, *p* = 2.34 *×* 10^−8^).

## Supplemental Video Legends

**Supplementary Video 1. Raw footage of bait-chasing behavior**. Black-and-white video recording of a representative active chasing bout. The animal pursues a moving bait attached to a manually operated fishing rod within the open arena. The bait trajectory was controlled by the experimenter to elicit sustained pursuit behavior while maintaining the bait within the animal’s visual field. Video is shown at real-time playback speed. NOTE: *Any human features visible in this video correspond to the main author of the study, who has provided consent for their inclusion*.

**Supplementary Video 2. Egocentric visualization of target-tuned spiking activity**. Animal-centered representation of chasing behavior, in which the animal’s position and heading are fixed at the center of the frame while the environment and bait position move relative to the animal. The bait location is indicated by a red dot. Audio represents spike events from a representative egocentric target cell (ETC), with each spike rendered as an audible click. Note that spiking activity increases when the bait is positioned to the right of the animal’s heading, at approximately 50°eccentricity, demonstrating egocentric tuning to target direction.

**Supplementary Video 3. Simultaneous Bayesian decoding of egocentric target and boundary directions**. Real-time visualization of population-level decoding during a representative chasing bout. The animal’s trajectory is shown in pink, with cardinal directions labeled. The red dot indicates the true bait position; the purple vector shows the decoded egocentric target direction. The blue vector indicates the true direction to the wall at any given time-point, while the green vector shows the decoded egocentric boundary direction. The black arrow denotes the animal’s head direction. Inset text displays instantaneous decoding accuracy, including true and decoded bait distance and wall distance. Decoded directions were computed from simultaneous RSC population activity, demonstrating that egocentric representations of targets and boundaries can be read out in parallel.

## Notes

### Competing Interest Statement

The authors have declared no competing interest.

### Summary of Updates

Revised figures and text, additional supplemental files

